# Temporal dynamics and biogeography of sympagic and planktonic autotrophic microbial eukaryotes during the under-ice Arctic bloom

**DOI:** 10.1101/2024.04.26.591324

**Authors:** Clarence Wei Hung Sim, Catherine Gérikas Ribeiro, Florence Le Gall, Ian Probert, Priscilla Gourvil, Connie Lovejoy, Daniel Vaulot, Adriana Lopes dos Santos

**Affiliations:** Asian School of the Environment, Nanyang Technological University, 50 Nanyang Avenue, 639798, Singapore; GEMA Center for Genomics, Ecology & Environment, Universidad Mayor, 8580745, Santiago, Chile; Sorbonne Université, CNRS, UMR7144, Team ECOMAP, Station Biologique de Roscoff, 29680, Roscoff, France; Sorbonne Université, CNRS, FR2424, Roscoff Culture Collection, Station Biologique de Roscoff, 29680, Roscoff, France; Département de Biologie, Institut de Biologie Intégrative et des Systèmes, Université Laval, Quebec, QC G1R1V6, Canada; Department of Biosciences, University of Oslo, PO Box 1066 Blindern, 0316, Oslo, Norway

**Keywords:** Under-ice bloom, Sympagic algae, Metabarcoding, 18S rRNA, Biogeography

## Abstract

Photosynthetic microbial eukaryotes play a pivotal role as primary producers in the Arctic Ocean, where seasonal blooms within and below the ice are crucial phenomena, contributing significantly to global primary production and biogeochemical cycling. In this study, we investigated the taxonomic composition of sympagic algae and phytoplankton communities during the Arctic under-ice spring bloom using metabarcoding of the 18S rRNA gene. Samples were obtained from three size fractions over a period of nearly three months at an ice camp deployed on landfast ice off the coast of Baffin Island as part of the Green Edge project. We classified the major sympagic and phytoplankton taxa found in this study into biogeographical categories using publicly available metabarcoding data from more than 2 800 oceanic and coastal marine samples. This study demonstrated the temporal succession of taxonomic groups during the development of the under-ice bloom, illustrated by an overall transition from polar to polar-temperate taxa, in particular in the smallest size fraction. Overlooked classes such as Pelagophyceae (undescribed Pelagomonadales clade A1 and the genus *Ankylochrysis*), Bolidophyceae (Parmales environmental clade 2), and Cryptophyceae (*Baffinella frigidus*) might play a greater role than anticipated within the pico-sized communities in and under the ice pack during the pre-bloom period. Finally, we emphasize the importance of microdiversity, taking the example of *B. frigidus* for which a new strain isolated during Green Edge represents an ice ecotype while the type strain is clearly linked to marine waters.

## Introduction

Autotrophic microbial eukaryotes (“microalgae”) are the major primary producers in the Arctic Ocean, dominating both pelagic (phytoplankton) and ice-associated (sympagic) primary production. Sympa gic production tends to be lower than phytoplankton production, accounting for 2-10% of total Arctic primary production (Lim et al. 2022; Watanabe et al. 2019). Ice-associated algae undergo a spring bloom that typically occurs a few months before that of the phytoplankton community, when light intensity increases and snow depth is sufficiently thin to allow light transmission through the ice (Selz et al. 2018). During the transition from ice cover to melt pond formation, when phytoplankton biomass under the ice is still low, sympagic communities serve as a rich food source for both ice-associated and early spring under-ice grazers, such as amphipods and calanoid copepods respectively (Kohlbach et al. 2016).

The initiation of the under-ice phytoplankton bloom (UIB) typically coincides with the termination of the sympagic algal bloom (Mundy et al. 2014). The stratification of the under-ice sea surface layer caused by the influx of freshwater from ice and snow melt reduces convective nutrient transport to the ice, inducing nutrient limitation (Oziel et al. 2019). The decrease of snow cover also leads to changes in the physical environment of the ice, resulting in photo-inhibition and brine flushing of ice algae (Selz et al. 2018). In the water, the increase in light resulting from melting snow sets the conditions for the development of the phytoplankton under-ice bloom (Matthes et al. 2020; Oziel et al. 2019).

The mosaic of Arctic marine environments (open water, sea ice, melt ponds, etc.), each with a wide range of nutrient and irradiance conditions, harbours complex heterogeneous communities of ice algae and phytoplankton (Ardyna et al. 2020a; Hop et al. 2020). Sympagic assemblages and bottom-ice communities tend to be dominated by pennate diatoms of the genera *Nitzschia*, *Fragilariopsis*, *Navicula* and *Cylindrotheca* (Ardyna et al. 2020a; Hop et al. 2020; Leu et al. 2015) and by the strand-forming centric diatom *Melosira* (Poulin et al. 2014). Centric diatoms such as *Thalassiosira* and *Chaetoceros* have been reported as major members of the phytoplankton community (Morando and Capone 2018; Oziel et al. 2019). Other photosynthetic groups also play a pivotal role in Arctic ecosystems. Three pico-sized (0.2-3 *µ*m) Mamiellophyceae genera, *Micromonas*, *Bathycoccus* and *Mantoniella*, can be abundant components of phytoplankton communities both under-ice and in ice-free waters (Balzano et al. 2012; Joli et al. 2017; Lovejoy et al. 2007; Not et al. 2005). The bloom-forming haptophyte *Phaeocystis pouchetii* has also been reported to dominate pelagic communities even under thick snow-covered pack ice (Assmy et al. 2017; Lasternas and Agustı 2010).

Major changes in the Arctic cryosphere are significantly altering the structure of sympagic and planktonic communities. The most emblematic and well documented impact of climate change in the Arctic is the rapid decline of sea ice cover in terms of area, thickness and age (Stroeve and Notz 2018). As less sea ice is retained during summer, multiyear ice has been replaced by younger and thinner first-year ice (Kwok 2018; Maslanik et al. 2011). Although first-year ice is more favourable to sympagic algae growth given its higher light transmittance (Lim et al. 2022), multiyear sea ice harbours a more diverse community of microbial eukaryotes (Hop et al. 2020). In the water, the northward flow of relatively warm North Atlantic water into the Arctic Ocean has not only amplified the decline of sea ice (Polyakov et al. 2017), but also triggered poleward intrusion of phytoplankton species of temperate origin in the European Arctic (Hegseth and Sundfjord 2008; Kwok 2018; Neukermans et al. 2018; Oziel et al. 2020). Warm anomalies in the Atlantic Water inflow to the Arctic Ocean have been linked to a shift from diatom-dominated phytoplankton communities to dominance by small coccolithophores (Lalande et al. 2013; Smyth et al. 2004).

The Arctic is anticipated to experience the highest species turnover in terms of invading and locally extinct species (Fossheim et al. 2015). Gains and losses of species in response to the ongoing changes in Arctic habitats (e.g., decrease in sea ice coverage and increased seawater temperature) are likely to induce significant food web reorganization with potential cascade effects (Beaugrand et al. 2019; Kortsch et al. 2015). Microbial eukaryotes differ in their thermal tolerance (Demory et al. 2019), dispersal capacity (Villarino et al. 2018) and ability to exploit new resources, (Andrew et al. 2023) which contributes to their abundance and diversity patterns. Therefore, they are natural proxies of community turnover and ecosystem shifts. A limited number of studies have contributed to establishing a baseline for the pan-Arctic microbial eukaryotic community of dinoflagellates (Okolodkov and Dodge 1996), diatoms (Supraha et al. 2022), mixotrophic (Stoecker and Lavrentyev 2018), and both autotrophic/non-autotrophic groups (Ibarbalz et al. 2023; Poulin et al. 2011). Two of these studies have dedicated specific efforts to linking taxonomically annotated 18S rRNA sequence “metabarcodes” from the Arctic to their biogeographical categorization (i.e. Arctic-temperate, cosmopolitan, etc), thus providing an overview of the biogeography of key Arctic phytoplankton taxa (Ibarbalz et al. 2023; Supraha et al. 2022).

The present study aimed to characterize Arctic sympagic algae and under-ice phytoplankton communities during the under-ice spring bloom using 18S rRNA gene metabarcoding. Samples were collected from three size fractions over nearly three months at a landfast ice camp off Baffin Island as part of the Green Edge project (Massicotte et al. 2020). The biogeographical distribution of major taxa identified during the bloom was analysed using over 2 800 marine samples from publicly available metabarcoding data. Our results describe the temporal succession of taxonomic groups during the development of the under-ice bloom, with a shift from polar to polar-temperate taxa, notably in the smallest size fraction. Additionally, it highlights the role of previously overlooked Arctic groups like Pelagomonadales and Parmales.

## Material and Methods

### Study area and sample collection

The field campaign was conducted on landfast sea ice, 360 m above the sea floor on the western coast of Baffin Bay (67.4797 *^◦^*N, 63.7895 *^◦^*W, Figure 1), as part of the Green Edge project. Sampling was carried out every two days between 4 May - 18 July 2016 at an ice camp set up southeast of Qikiqtarjuaq Island (Nunavut). Bottom-ice was collected from two sections of ice cores: (i) 0-3 cm (ICE_0) and (ii) 3-10 cm (ICE_1) above the bottom of the core (Figure 2). Ice slices were melted in 0.2 *µ*m filtered seawater. Three litres of under-ice water were collected using Niskin bottles at four depths: (i) 1.5 m (WATER_1), (ii) 5-10 m (WATER_2), (iii) 10-20 m (WATER_3) and (iv) 40-60 m (WATER_4) (Figure 2). Samples were pre-filtered with a 100 *µ*m mesh to remove metazoans and then size fractionated through a peristaltic line with two 47 mm Swinnex holders with 20 *µ*m and 3 *µ*m polycarbonate filters, and a 0.2 *µ*m Sterivex at the end of the line, to obtain micro (20-100 *µ*m), nano (3-20 *µ*m), and pico (0.2-3 *µ*m) size fractions. Polycarbonate filters were folded carefully and preserved with 1.8 mL of RNALater^®^. All samples were stored at -80*^◦^*C until processing.

**Figure 1:**
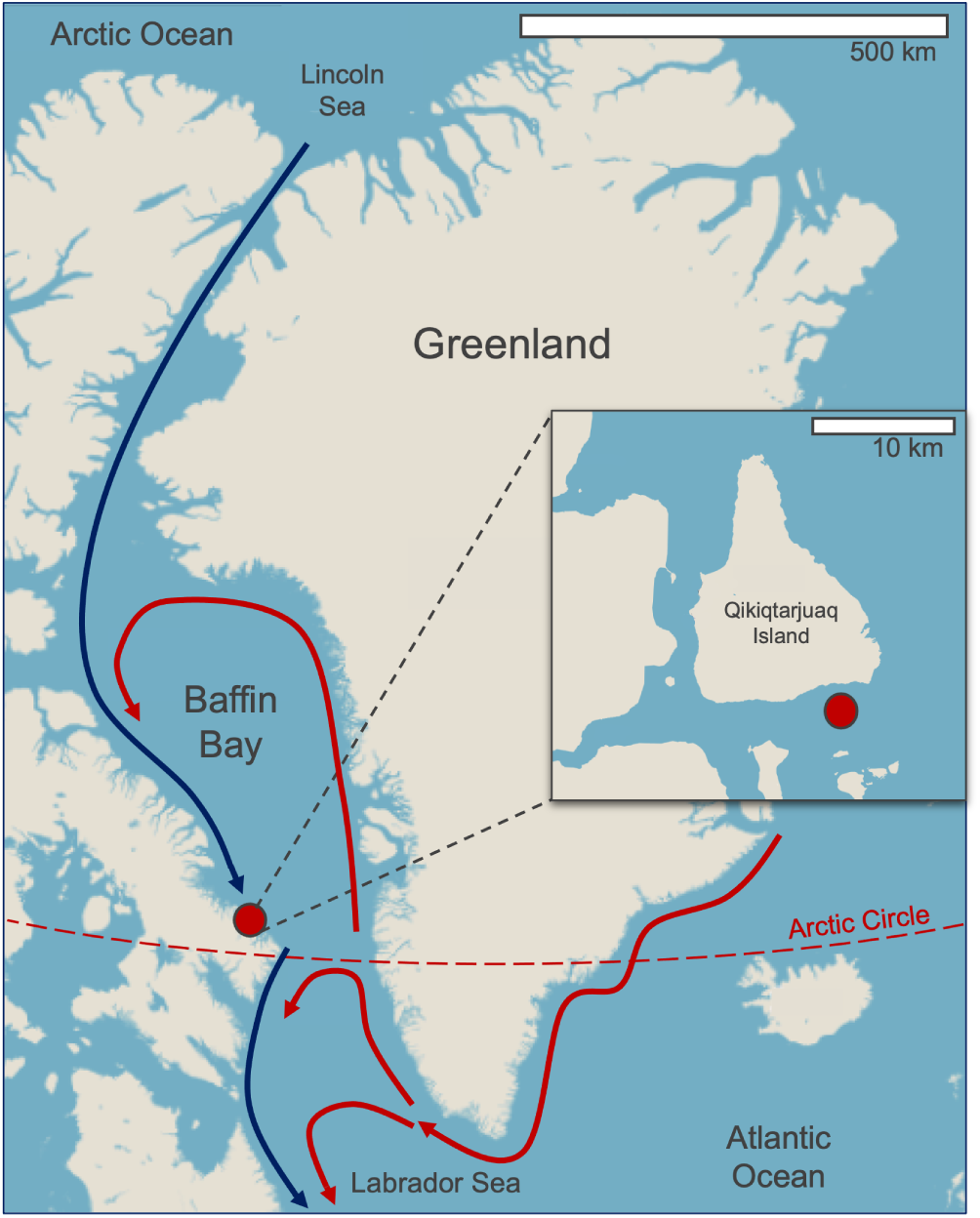
Location of the ice camp on landfast sea ice near Qikiqtarjuaq Island on the western coast of Baffin Bay. Arrows indicate a simplification of the counter-clockwise Atlantic-derived (red) and Arctic-derived (blue) water mass circulation, adapted from Tang et al. (2004).

**Figure 2:**
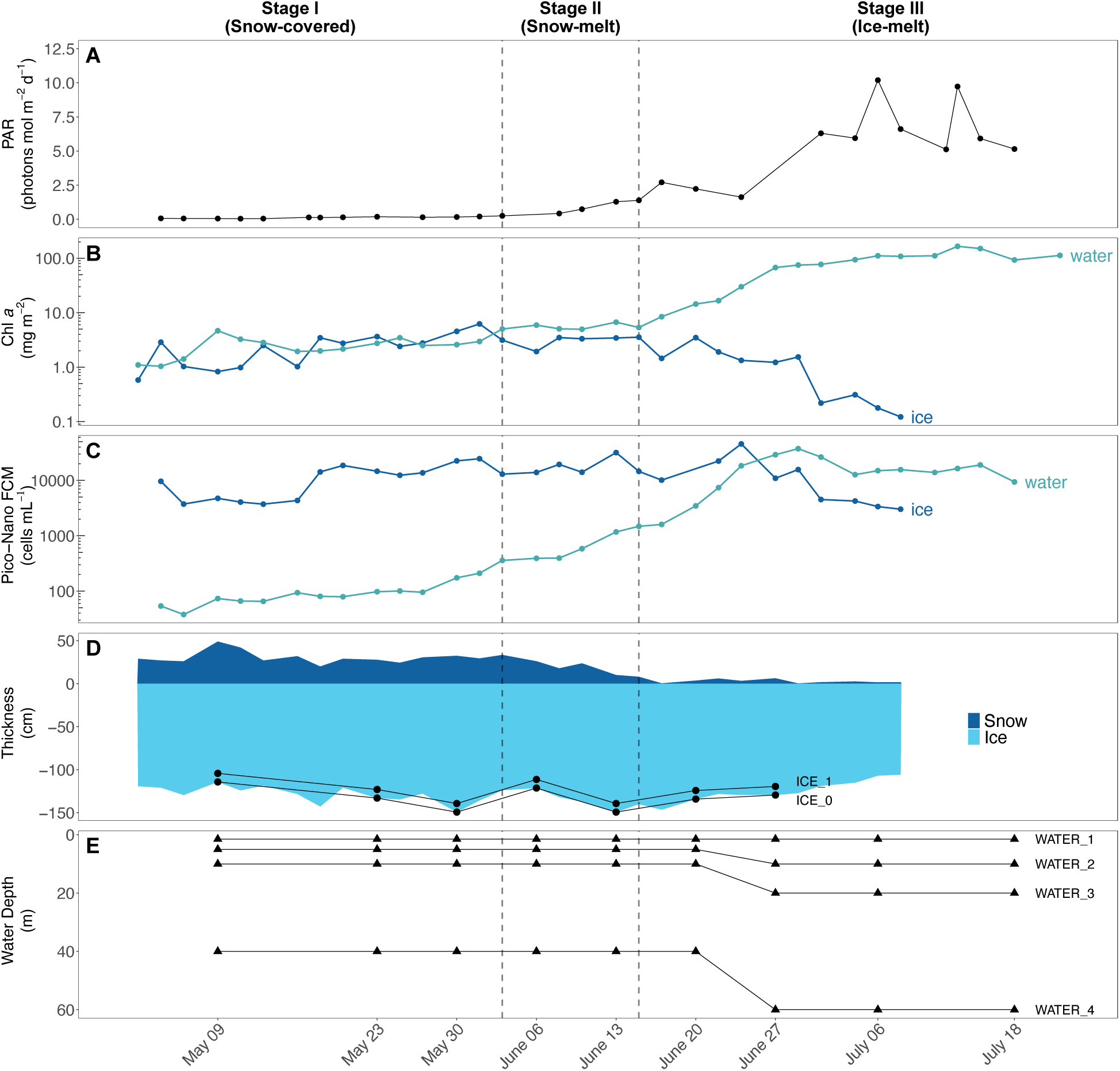
Environmental variables and sampling through the three main stages of the under-ice phytoplankton bloom. **(A)** Under-ice PAR at 2 m depth (mol photons. m*^−^*^2^. d*^−^*^1^); **(B)** Surface integrated Chl *a* concentrations (mg. m*^−^*^2^) from ice (bottom 10 cm) and water column (top 60 m); **(C)** Pico/nano—phytoplankton abundance (cells. mL *^−^*^1^); **(D)** Ice and snow thickness (cm). ICE_0 and ICE_1 represent samples from the bottom 3 cm and 3-10 cm of ice; **(E)** Water samples collected at four levels. No data were available for ice and snow thickness after July 08.

### Environmental data

Photosynthetically active radiation (PAR) was computed from 19 discrete spectral irradiance wave-lengths (380-875 nm) measured using an ICE-Pro (an ice flow version of the Compact-Optical Profiling System, C-OPS). Sea ice and under-ice water Chlorophyll *a* (Chl *a*) concentrations were obtained by high-performance liquid chromatography (HPLC). Water Chl *a* concentration (mg m*^−^*^2^) was depth-integrated from four discrete depths corresponding to water samples obtained in the first 60 m of the water column, while depth-integrated ice Chl *a* (mg m*^−^*^2^) was derived from the bottom 10 cm of the ice. Pico- and nano-phytoplankton cell abundance was measured using a BD Accuri*^T M^* C6 flow cytometer as previously described in Massicotte et al. (2020). All ancillary physico-chemical and biological data obtained from the Green Edge project are available at SEANOE. Variables characterizing the environmental conditions during the time series such as snow and ice thickness, under-water PAR, Chl *a* biomass and photosynthetic pico-nano cell abundances were used to group the samples into the three bloom stages described by Ardyna et al. (2020b): (I) snow-covered, (II) snow-melt and (III) ice-melt (Table 1, Figure 2).

**Table 1:** Stages of the Arctic spring bloom in Baffin Bay 2016 ice-camp following Ardyna et al. (2020b)

| Stage no. | UIB stage | Dates |
| --- | --- | --- |
| Stage I | Snow-covered | 01 May - 03 June 2016 |
| Stage II | Snow-melt | 03 June - 15 June 2016 |
| Stage III | Ice-melt | 15 June - 18 July 2016 |

### DNA extraction, 18S rRNA V4 gene PCR amplification and sequencing

Samples for DNA extraction were selected from each stage of bloom based on the evolution of Chl *a* and autotrophic cell abundance determined by flow cytometry (Figures 2B, 2C). DNA was extracted using ZR Fungal/Bacterial DNA MiniPrep (Zymo Research, Irvine, CA, USA) following the manufacturer’s instructions, and final DNA concentration was measured using PicoGreen™(Thermo Fisher Scientific, Waltham, MA, USA). The 18S rRNA V4 hypervariable gene region (around 380 bp) was amplified with the primers TAReuk454FWD1 (forward, 5’-CCAGCASCYGCGGTAATTCC-3’) and V4 18S Next.Rev (reverse, 5’-ACTTTCGTTCTTGATYRATGA-3’) (Piredda et al. 2017). Reaction mixtures (20 *µ*L) were performed using 10 *µ*L of Phusion High-Fidelity PCR Master Mix^®^ 2*×*, 0.3 *µ*M final concentration of each primer, 3% DMSO, 2% BSA, 5 ng of template DNA and H_2_O. Thermal conditions were as follows: 98*^◦^*C for 5 min, followed by 25 cycles of 98*^◦^*C for 20 s, 52*^◦^*C for 30 s, 72*^◦^*C for 90 s, and a final cycle of 72*^◦^*C for 5 min. Samples were amplified in triplicate and pooled together. PCR purification, library preparation and amplicon sequencing was conducted at the GeT-PlaGe platform of GenoToul (INRAE Auzeville, France) using an Illumina Miseq and the 2 x 250 cycles Miseq kit version 2.

### Sequence processing, trophic mode allocation and culturability

Sequences were processed with scripts written in the R language (R Core Team 2020) using the *dada2* package (Callahan et al. 2016). Primer sequences were first removed with *cutadapt* version 2.8 (Martin 2011) using the default parameters. Reads were filtered and trimmed using the filterAndTrim function with the following parameters: truncLen = c(230, 230), maxN = 0, maxEE = c(2, 2), and truncQ=10. Forward and reverse reads were merged with the mergePairs function and chimeric sequences removed with the removeBimeraDenovo function, both using default parameters. Amplicon Sequence Variants (ASVs) obtained with the dada2 function were labelled with the first 10 characters of the 40 character hash value of the sequence computed using the sha1 function from the R *digest* package (Eddelbuettel 2021). ASVs were taxonomically assigned using assignTaxonomy function with PR^2^ database version 5.0.1 (https://pr2-database.org, Guillou et al. 2012) as a reference. ASVs with low (*<* 80%) bootstrap support from the assignTaxonomy function at a given taxonomic level were reclassified to the next higher taxonomic level until the bootstrap value was *≥* 80%. ASVs assigned to non-protist taxa (e.g. metazoans) were removed. This included all taxa from divisions Metazoa, Fungi, Rhodophyta, classes Phaeophyceae, Embryophyceae, orders Bryopsidales, Ulotrichales, Dasycladales, Trentepohliales, Cladophorales and unidentified Opisthokonta. This resulted in a total of 2196 ASVs.

ASVs were then assigned to a trophic mode (photosynthetic, mixotrophic, heterotrophic, dinoflagellate) based on the database from Schneider et al. (2020). Only ASVs classified as photosynthetic and mixotrophic were further analysed in this study. Non-constitutive mixoplankton that do not have the innate ability to perform photosynthesis were also not considered, resulting in the removal of Ciliophora and Rhizaria. Finally, dinoflagellates were not considered, since this group contains taxa that have a range of trophic modes, which can vary even within a given genus (Cohen et al. 2021). ASVs corresponding to taxa not present in the Schneider database were allocated to a trophic mode using a majority rule. For example, within the class Prymnesiophyceae, 15 out of the 23 listed taxa were assigned as mixotrophic and the other 8 as photosynthetic. All other Prymnesiophyceae taxa not present in the Schneider database were then assigned as mixotrophic. The majority rule was only applied to groups that have at least 5 taxa in the Schneider database. A few higher level taxa that were not present in the Schneider data were assigned to a trophic mode based on the literature. A total of 428 ASVs were classified as photosynthetic and mixotrophic and further considered in this study. The similarity of ASVs to sequences of taxa available in cultures was determined by the – usearch_global option of *vsearch* with iddef = 2 against culture sequences from the PR^2^ database version 5.0.1 (Guillou et al. 2012). Finally, the number of reads in each sample was normalized by the median dataset sequencing depth (10 231 reads).

### Biogeographical distribution of ASVs

To determine the biogeographical distribution of ASVs, we used version 2.0 of the metaPR^2^ database (Vaulot et al. 2022), which contains 18S rRNA metabarcodes from published studies re-processed with dada2 and annotated with the PR^2^ reference sequence database. We used a total of 2 874 marine samples (oceanic and coastal) using the V4 region 18S rRNA gene (Table S1). Samples were classified as polar (*≥* 66 *^◦^*N and *≥* 66 *^◦^*S), temperate (23-66 *^◦^*N and 23-66 *^◦^*S) and tropical (23 *^◦^*S - 23 *^◦^*N) (Figure S1).

ASVs from the present study and those from metaPR^2^ were clustered (cASVs) if they showed 100% similarity in their overlap regions (see Vaulot et al. 2022). cASVs were then assigned to a biogeographical category (e.g. polar, temperate, cosmopolitan), based on their occurrence in metaPR^2^ samples following the approach of Supraha et al. (2022) (Table 2). cASVs were assigned to a biogeography category if at least 90% of the samples where they occurred fell within the regions defined in Table 2. cASVs were assigned as cosmopolitan if they were present in all three major regions (polar, temperate, tropical) but did not have a clear dominance in any region. cASVs that were present in less than five samples were considered unallocated. Multiple samples from a single geographical point (in particular from time series studies) were considered as a single occurrence. In the rest of the paper, we refer to cASV as ASV for simplicity.

**Table 2:** Criteria for biogeography classification of ASVs presented in this study based on their occurrence in 2,874 samples from public available datasets gathered in metaPR^2^.

| Biogeography | Description | ASV occurrence |
| --- | --- | --- |
| Polar | ASV mostly restricted to the Arctic and Antarctic | Polar ≥ 90% |
| Polar-Temperate | ASV present in the polar and temperate regions | Polar + Temperate ≥ 90% |
| Temperate | ASV mostly restricted to the temperate region | Temperate ≥ 90% |
| Temperate-Tropical | ASV present in the temperate and tropical regions | Temperate + Tropical ≥ 90% |
| Tropical | ASV mostly restricted to the tropical region | Tropical ≥ 90% |
| Cosmopolitan | ASV has a global distribution | Polar, Temperate , Tropical $>$ 0% each |
| Unallocated | ASV is either unique to this study or has been found in less than five metaPR<sup>2</sup> samples |  |

### Data analysis and visualization

Data analysis was performed within R, using the following packages: *tidyr* (Wickham 2021) and *dplyr* (Wickham et al. 2021) for filtering and organizing data; *ggplot2* (Wickham 2016) for data visualization; *treemapify* (Wilkins 2021) for treemaps, *ggridges* (Wilke 2021) for density plots, *ggcharts* (Neitmann 2021) for lollipop charts, and *patchwork* (Pedersen 2020) for merging plots. Non-metric multidimensional scaling (NMDS) ordination of a Bray-Curtis dissimilarity matrix was performed with *phyloseq* (McMurdie and Holmes 2013). ANOSIM (ANalysis Of SIMilarity) (package *vegan* Oksanen et al. 2020) was used to assess the influence of size fraction, substrate (ice and water) and bloom phase on the community composition. Indicator species analysis (*indicspecies* package, (De Caceres and Legendre 2009)) was performed on ASVs within each size fraction in order to find significant association between taxa and substrate (ice vs. water) and bloom stages (stage I vs. stages II and III). Default *IndVal* index was used as a statistical test with 9 999 random permutations. The following color-vision-deficiency friendly palettes were used: BrBg, Blues, Reds and Set1 from *RColorBrewer* (Neuwirth 2014) and Okabe-Ito from base R (R Core Team 2020).

### Scanning electron microscopy

Ice (100 mL) and water (200 mL) samples were filtered through 0.8 *µ*m size polycarbonate (Isopore or Nuclepore) membranes using a vacuum pump (< 250 mm Hg) for scanning electron microscopy. Samples were left to dry in an oven at 35 *^◦^*C for 1 hour and subsequently stored at room temperature. Filters were cut, mounted on stubs with carbon adhesive paper, metallized with palladium gold and observed with a Scanning Electronic Microscope (SEM, Phenom G2Pro, PhenomWorld) at 10 kV.

## Results

The Green Edge Ice Camp took place from spring to early summer, spanning periods of ice-covered to ice-free waters. The study site was located on landfast sea ice on the western side of Baffin Bay south of Qikiqtarjuaq Island (Figure 1). Baffin Bay is a seasonally ice-covered regional sea within the Canadian Arctic. As ice melts in spring, the sea ice edge retreats westwards from Greenland towards Canada. Water masses in Baffin Bay circulate counter-clockwise. Warm and salty Atlantic-derived waters enter the Bay through the Davis Strait, north of the Labrador Sea, and move northwards along the coast of Greenland (Tang et al. 2004). Cold Arctic-derived waters entering northern Baffin Bay move southwards along the coast of Canada, eventually flowing out of southwestern Baffin Bay (Münchow et al. 2015) (Figure 1).

PAR at 2 m (ice-water interface), used as a proxy of the evolution of under-ice solar radiation available for photosynthesis, remained consistently low during snow-covered stage I (Table 1, with values between 0.07-0.25 mol m*^−^*^2^ d*^−^*^1^ until mid-June (Figure 2A). Concomitant with the drop in surface albedo of snow, PAR increased from 2.7 mol m*^−^*^2^ d*^−^*^1^ to 10.2 mol m*^−^*^2^ d*^−^*^1^ between stages II and III. Bottom-ice Chl *a* and sympagic photosynthetic pico-nano (0.2-20 *µ*m) cell concentration remained stable until the end of the snow-melt period. Bottom-ice Chl *a* concentration peaked at 6.2 mg m*^−^*^2^ in early June, while photosynthetic pico-nano (0.2-20 *µ*m) cell concentration reached its maximum value (35,000 cell mL*^−^*^1^) by the end of June. The sympagic photosynthetic community then slowly declined towards the end of the sampling period during stage III (Figures 2B, 2C).

Under-ice Chl *a*concentration and pico-nano phytoplankton cell abundance remained consistently low from May to mid-June (Figures 2B, 2C). Both parameters peaked (182 mg m*^−^*^2^ and 45,000 cells mL*^−^*^1^) during the first week of July, when the absence of snow and the presence of melt ponds allowed the increase of light (PAR peaked at 10.2 *µ*mol m*^−^*^2^, Figure 2A), setting the conditions for the development of the under-ice phytoplankton bloom.

### Community diversity and structure

Sympagic and under-ice planktonic communities of microbial eukaryotes were separated according to size fractions (0.2-3 *µ*m, 3-20 *µ*m and > 20 *µ*m) and their diversity was accessed by metabarcoding of the 18S rRNA gene. Four hundred and twenty-eight ASVs were assigned to photosynthetic taxa. The ice algal and phytoplankton communities were composed of 18 and 21 classes, respectively, including Bacillariophyceae, Mediophyceae, Pelagophyceae, Chrysophyceae and Bolidophyceae from the division Stramenopiles, Mamiellophyceae, Pyramimonadophyceae and Chlorophyceae from the Chlorophyta, Prymnesiophyceae (Haptophyta) and Cryptophyceae (Cryptophyta) (Figure S2). Morphological identification of Bacillariophyceae and Mediophyceae taxa from both sympagic (Figure S3) and phytoplanktonic communities (Figures S4 and S5) was used to complement the metabarcoding data. Few genera and species were identified by both methods (Tables S2, S3). Twenty species were uniquely identified by SEM for ice (9) and water (11) samples (Tables S2, S3). The majority of ASVs obtained in this study had no match to public sequences from cultures (Figure S6). Community analysis at ASV level using NMDS and ANOSIM showed that samples clustered according to size fractions along the first axis and substrate (ice and water) along the second axis (Table S4, Figure S7). Community composition was significantly different between the three stages of UIB for both ice (*R* = 0.15; *p* = 0.004) and water (*R* = 0.420, *p* = 0.001) (Table S4). Twenty-three key taxa accounting for 75% of the total photosynthetic reads were selected to analyze community change across bloom stages and size fractions (Figure 3).

**Figure 3:**
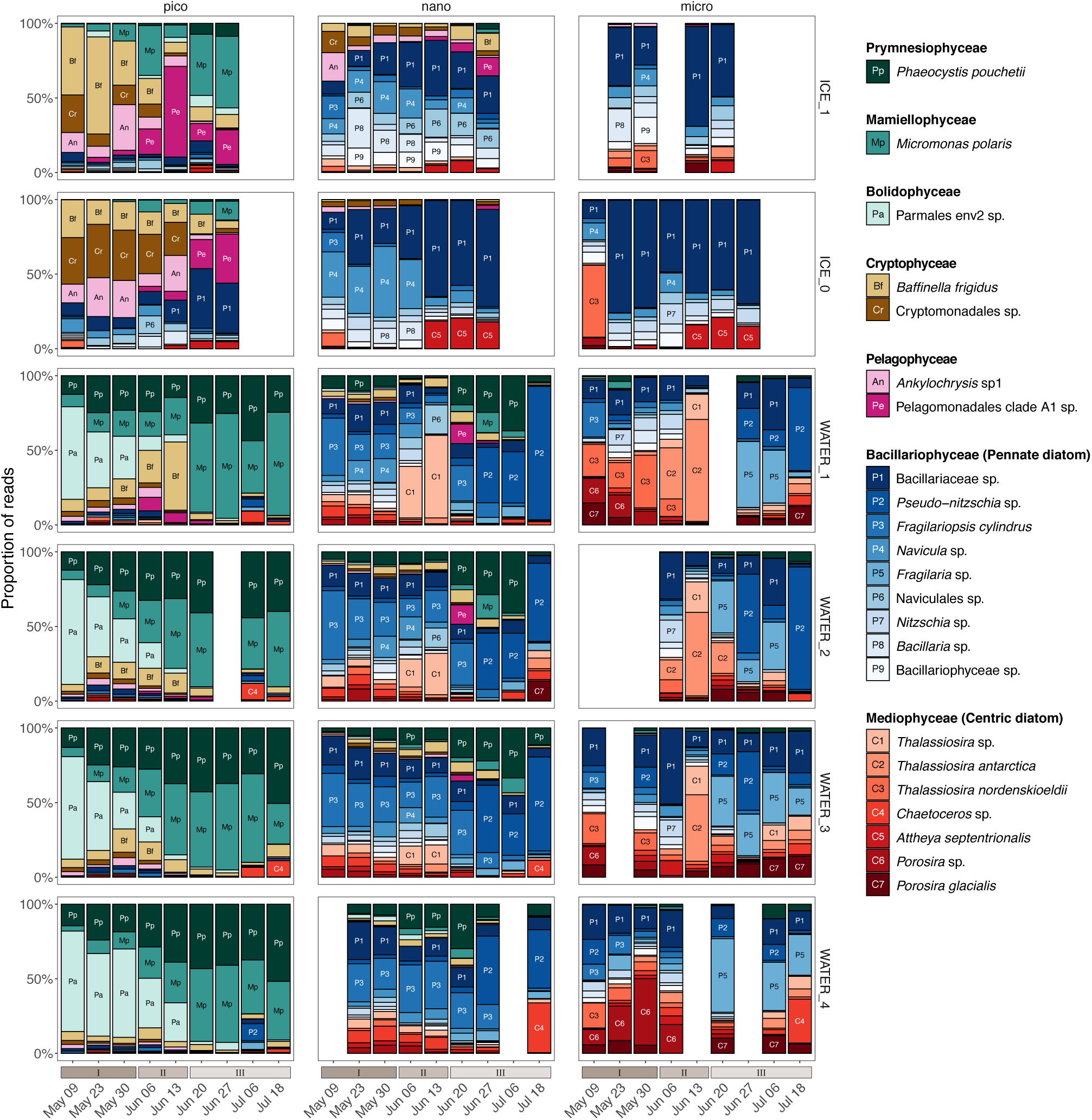
Relative abundance of 18S V4 rRNA reads at the species level from 23 most abundant photosynthetic taxa across bloom stages I to III in ice and water. These species represent 75% of all reads. Samples are sorted according to their size fractions: micro (20-100 *µ*m), nano (3-20 *µ*m) and pico (0.2-3 *µ*m). Taxa with a minimum of 10% relative abundance of samples are annotated with a two character label. No ice samples were collected in July when melt ponds were formed and the ice was instable. Other blank columns represent samples lost during processing.

Within the sympagic pico-sized community, during the snow and ice covered stages (I and II) the Cryptophyceae *Baffinella frigidus* and an undescribed Cryptomonadales clade were dominant in the 3-10 cm ice layer (ICE_1) and bottom 3 cm (ICE_0), respectively. At the same time, in (ICE_0), *Ankylochrysis* (Pelagophyceae) and the undescribed Pelagomonadales clade A1 co-dominated in the bottom ice layer. During stage III, the Mamiellophyceae (*Micromonas polaris*) was abundant in the 3-10 cm ice layer (ICE_1) (Figure 3). Among pico-sized phytoplankton in water, the dominant taxon during stage I belonged to Parmales environmental clade 2 with a contribution increasing with depth (Figure 3). From stage II onwards, it was replaced by *M. polaris* and the haptophyte *P. pouchetii*. In the surface layer (WATER_1), the cryptophyte *B. frigidus*, which was present in the ice earlier, was abundant during stage II.

The nano-sized communities, both in the water and ice, were dominated in terms of abundance and diversity by Bacillariophyceae (pennate diatoms). In the ice, during stage I, a diverse community of nano-sized pennate diatoms inhabited the 3-10 cm layer (ICE_1) without clear dominance of any genus or clade (Figure 3). Within the bottom layer (ICE_0), *Navicula* and unassigned Bacillariaceae ASVs co-dominated the community (Figure 3) until the beginning of stage II. From this point, several ASVs assigned to undescribed pennate diatoms (Bacillariaceae) co-dominated the nano-sized sympagic community until the end of the sampling period. There was also a small contribution to the sympagic community from the centric diatom *Attheya septentrionalis* during the transition from stage II to III (Figure 3). A pennate-centric-pennate diatom temporal succession was observed for the nanoplankton in the surface water layer, featuring the genera *Fragilariopsis* (*F. cylindrus*), *Thalassiosira* and *Pseudo-nitzschia*.The haptophyte *P. pouchetii* also contributed significantly to the nano-sized community in the water column during stage III.

The micro-sized ice community from stages I and II was also dominated by undescribed Bacillariaceae diatoms, except for bottom ice at the start of stage I when *Thalassiosira nordenskioeldii* was the dominant taxon (Figure 3). Microplanktonic diatoms were co-dominated by genera and undescribed groups of pennate (Bacillariophyceae) and centric (Mediophyceae) diatoms (Figure 3). The centric diatom genus *Thalassiosira* (*T. antarctica and T. nordenskioeldii*) increased its contribution in the surface layer (WATER_1) during stage II, while *Porosira* had a high contribution to the deep water layer community during stage I (Figure 3). Among pennate diatoms, *Pseudo-nitzschia*, *Fragilaria* and undescribed Bacillariaceae dominated the microplanktonic community during stage II without a clear pattern in abundance (Figure 3).

### Biogeography and microdiversity

We explored the occurrence of our ASVs across 2,874 marine samples selected from the metaPR^2^ database (Figure S1, Table S1) and classified them according to their latitudinal occurrence (Table 2). A total of 200 ASVs, representing 82.5% of photosynthetic reads, were assigned to a biogeographical region (Figure 4). Most of the allocated ASVs had polar and polar-temperate distributions, with few temperate and cosmopolitan ASVs (Figure 4). The sympagic communities were dominated by polar ASVs in all size fractions, and also harbored most of the unallocated ASVs. The pico-phytoplankton community was initially co-dominated by polar and polar-temperate ASVs (Figure S8), but as the bloom developed, ASVs assigned as polar-temperate represented most of the reads. In contrast, the nano fraction was dominated by polar-temperate ASVs and no clear change was observed. Finally, among the micro-sized phytoplankton, polar ASVs were important at almost all times, except during the last sampling week (stage III) in surface waters (WATER_1 and 2), where polar-temperate ASVs became dominant (Figure S8).

**Figure 4:**
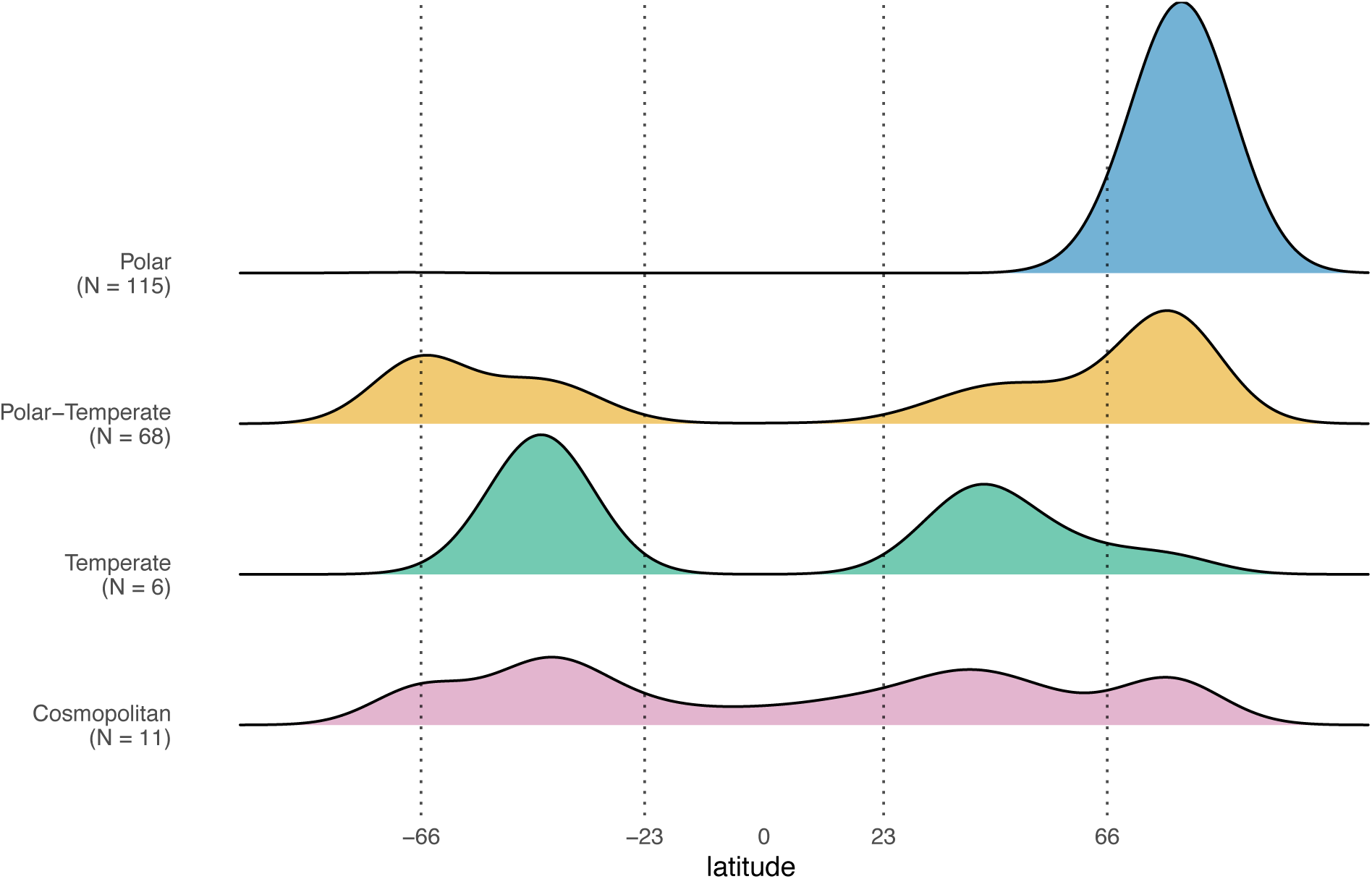
Latitudinal distribution of the metaPR^2^ samples where Green Edge ASVs occurred. Polar, Polar-temperate, Temperate and Cosmopolitan refer to the biogeographic classification of the ASVs (Table 2. *N* indicates the number of assigned ASVs in each category. The major latitudinal boundaries are illustrated by dashed lines: Arctic circle (66 *^◦^*N), Tropic of Cancer (23 *^◦^*N), Tropic of Capricorn (23 *^◦^*S) and Antarctic circle (66 *^◦^*S).

We also investigated whether the 20 most abundant ASVs for each substrate and bloom stage were significantly associated with a given substrate (ice vs. water) or bloom stage using indicator species analysis. The samples from stage I were grouped as dark phase and stages II and III as light phase, based on the recorded PAR (Figure 2A), to facilitate pairwise comparison (Figure 5).

**Figure 5:**
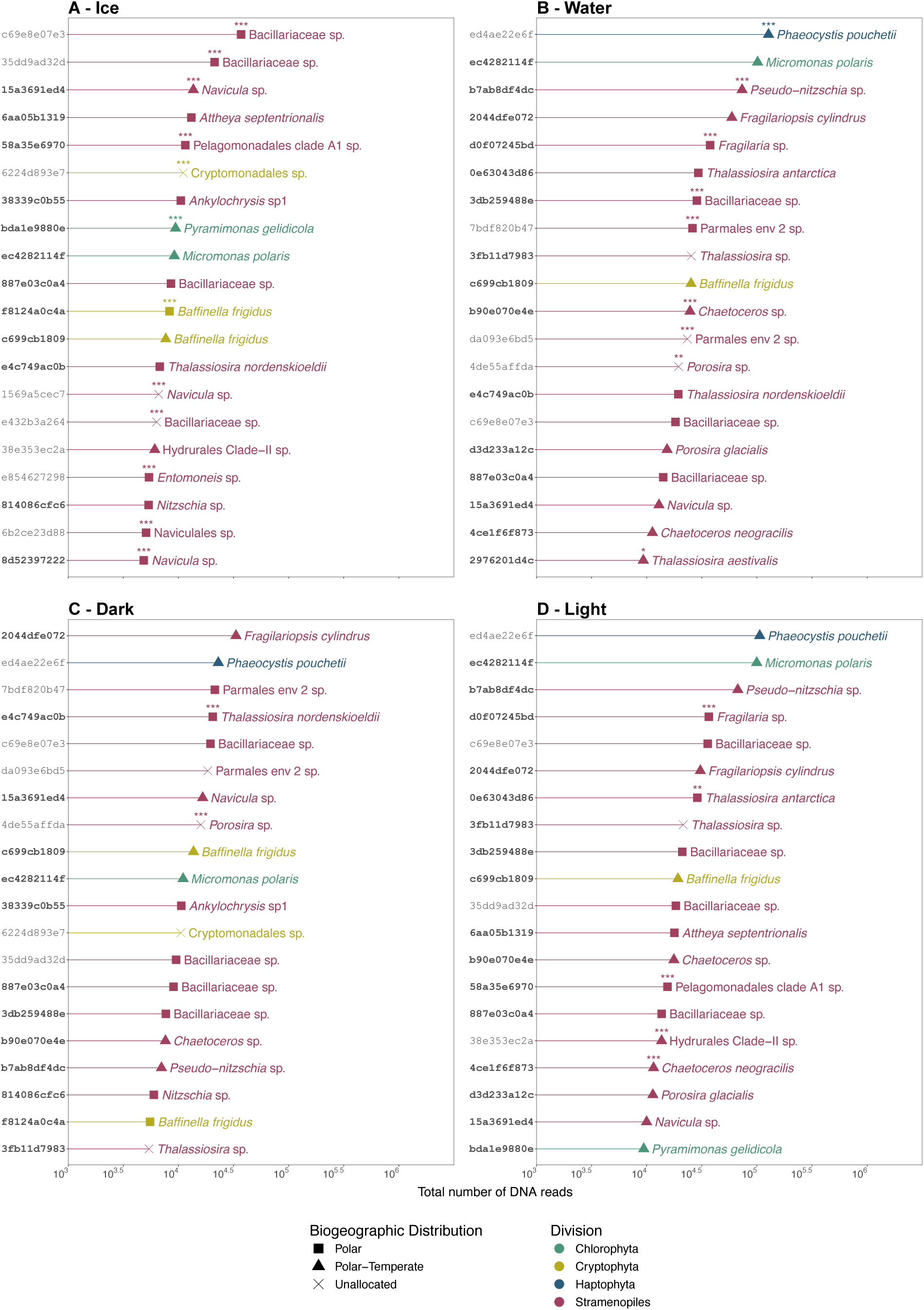
Twenty most abundant photosynthetic ASVs in ice (A) and water (B) substrates, and UIB dark (stage I, C) and light (stages II+III, D) phases. Symbol shape corresponds to biogeographical classification. ASVs in bold correspond to those whose sequence matches 100% to PR^2^ sequences obtained from cultures. Colours correspond to division. The *indicspecies* statistical significance is shown as follows: *p < 0.05, **p < 0.01, ***p < 0.001.

Twelve and 9 ASVs, respectively, were significantly associated with ice and water. Most of the ice associated ASVs (7) were allocated as polar (Figure 5A), while those flagged as indicators of the water community were equally split as polar, polar-temperate and unallocated (Figure 5B). Fifty percent of the ASVs flagged as indicators of ice and water substrates had a 100% match to at least one culture sequence (Figures 5A, 5B). Only 2 ASVs were flagged as indicators for the dark phase (*T. nordenskioeldii* ASV_e4c749ac0b and *Porosira* sp. ASV_4de55affda) (Figure 5C) and 5 ASVs were significantly associated with the light phase (Figure 5D). Among these, 85% had a 100% match to at least one culture sequence (Figures 5C, 5D). ASVs flagged as indicators from the genera *Baffinella* and *Thalassiosira* that exhibit diversity at the species or ASV level (Figure 5) were further investigated (Figure 6A).

**Figure 6:**
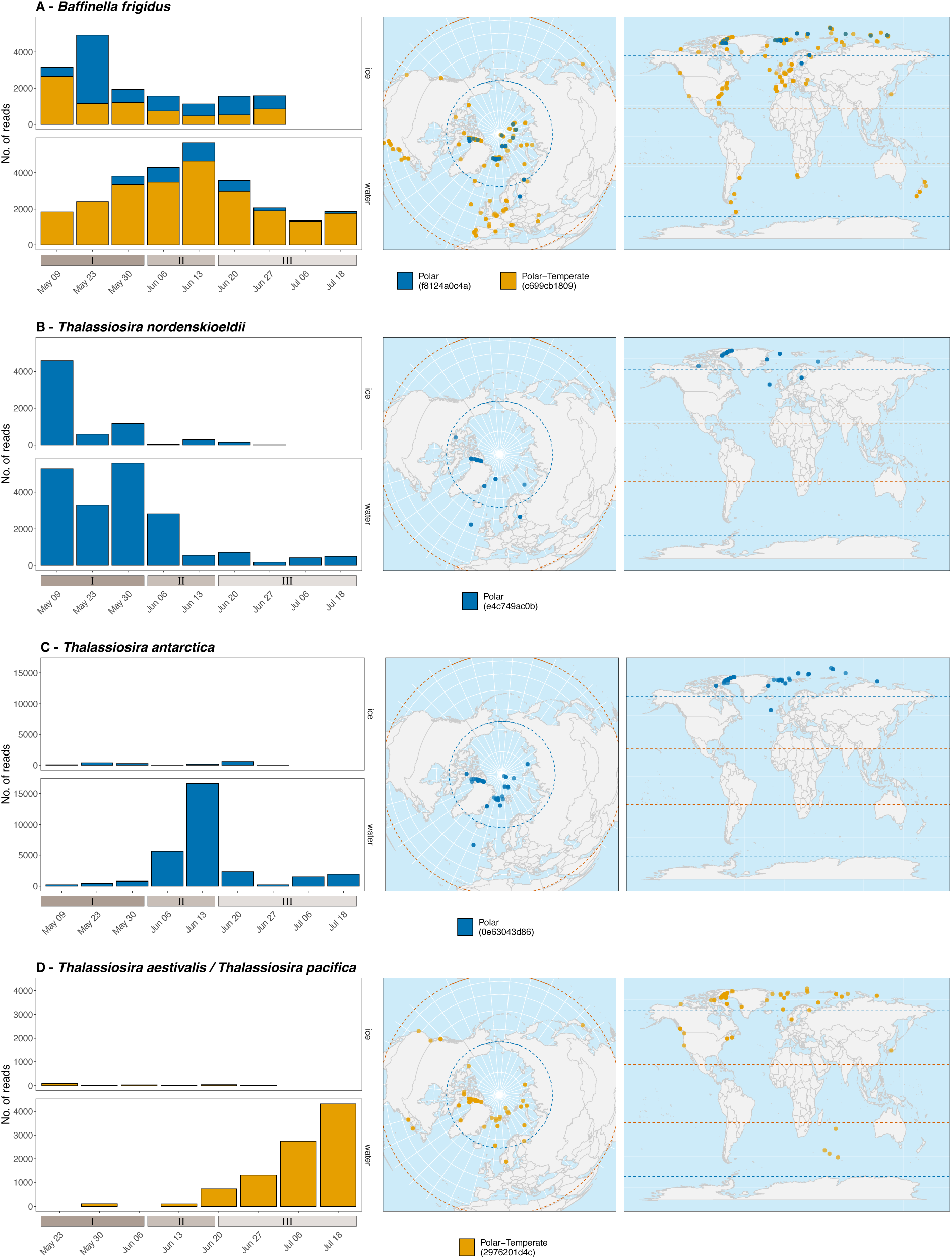
Temporal evolution and geographical occurrence of ASVs assigned to (A) *B. frigidus*, (B) *T. nor-denskioeldii*, (C) *T. antarctica* and (D) *T. aestivalis/T. pacifica*. Left: ASVs read numbers across different bloom stages in the ice and water samples (all depths and size fractions combined). Center and right: metaPR^2^ samples where the ASVs were detected. Dashed lines correspond to the polar circles and the tropics.

*B. frigidus* ASV_c699cb1809 (polar-temperate) and ASV_f8124a0c4a (polar) were co-dominant in the ice samples, whilst in the water samples the polar-temperate ASV_c699cb1809 was dominant (Figure6A). The polar ASV_f8124a0c4a was flagged as an ice indicator (*p* < 0.001, Figure 5A). While the polar-temperate ASV_c699cb1809 was more abundant in water samples (Figure 6A), this ASV was not significantly associated with either substrate (Figure 5). The contribution of the ice and polar ASV_f8124a0c4a increased in the water column during the transition between stages II and III (Figure 6A), possibly due to the release of cells from the ice during melting. One base pair differentiates these two *B. frigidus* ASVs (Figure S9).

Three *Thalassiosira* ASVs, flagged as indicators and with distinct biogeographical allocation, exhibited a succession pattern during the UIB (Figures 6B-D). *T. nordenskioeldii* ASV_e4c749ac0b and *T. antarctica* ASV_0e63043d86, both categorized as polar, were flagged as dark and light bloom phase indicators, respectively (Figures 5C, 5D), while the polar-temperate *T. aestivalis/T. pacifica* ASV_2976201d4c (*T. pacifica*/*T. aestivalis* share an identical V4 sequence, so both species names were used to refer to this ASV) was associated with water (Figures 5B). Based on read numbers, *T. nordenskioeldii* ASV_e4c749ac0b was more abundant in the water during the dark phase and then was replaced by *T. antarctica* ASV_0e63043d86 at the start of light phase (Figures 6C, 6D). Finally, the polar-temperate *T. aestivalis/T.pacifica* ASV_2976201d4c became the dominant *Thalassiosira* ASV in the water during the development of the UIB (Figure 6D).

## Discussion

### Taxa succession and diversity

The Green Edge campaign documented the taxonomic succession of sympagic and under-ice phytoplankton communities in the western Baffin Bay. During the snow-covered and snow-melt stages, photosynthetic biomass, both in and beneath the landfast sea ice, was limited by light availability, but still present as indicated by Chl *a* and photosynthetic pico-nano (0.2-20 *µ*m) cell concentrations (Figure 2). The melting of the snow allowed more light to reach the sub-surface water, inducing an increase in biomass and number of cells of the under-ice (pico and nano) phytoplankton community (Figure 2). Similar to other time series studies under landfast ice (Mundy et al. 2014), light availability and upper water column stabilization were the key factors triggering the under-ice bloom during the Green Edge campaign (Oziel et al. 2019). A similar trend was also observed during a campaign in the marginal ice zone (Randelhoff et al. 2019; Ribeiro et al. 2022).

Community analysis at the ASV level demonstrated that the sympagic and under-ice planktonic communities were significantly different between the three stages of the UIB (Table S4). Light is an important driver of taxonomic diversification. Sympagic and under-ice phytoplanktonic communities have evolved distinct strategies to exploit their dynamic light environment and to maximize light absorption. Pennate diatoms from sympagic and planktonic communities are important members of under-ice bloom and low-light dwellers (Hancke et al. 2018). In the present study, pennate diatoms, in particular raphid pennate genera such as *Navicula*, *Nitzschia*, *Pseudo-nitzschia*, and ribbon forming pennates *Fragilariopsis* and *Fragilaria*, were more diverse than the other autotrophic classes and dominated both sympagic and planktonic nano/micro-sized communities in terms of relative abundance through time (Figure 3). These taxa have been commonly reported in bottom-ice samples (Hegseth and von Quillfeldt 2022; Hop et al. 2020; Poulin et al. 2011). In contrast, the increased contribution of the epiphytic centric genus *Attheya*, concomitant with snow melt (Figure 2, Figure 3) was consistent with studies reporting the appearance of the centric diatom *A. septentrionalis* in bottom-ice communities as light intensities increase through spring (Campbell et al. 2018; Melnikov et al. 2002; van Leeuwe et al. 2022).

Arctic diatom assemblages are often described by their seasonal succession pattern from pennate to centric species linked to different light requirements (Ardyna and Arrigo 2020; Ardyna et al. 2020a). Among nanoplanktonic diatoms, a pennate-centric-pennate succession was observed in the surface water layer, where the relative abundance of centric diatoms represented by the genus *Thalassiosira* increased during stage II and receded during stage III (Figure 3). The succession observed at the ice camp was however not observed in under-ice samples collected during the Green Edge oceanographic campaign (Ribeiro et al. 2022), which occurred simultaneously offshore throughout the central Baffin Bay (Bruyant et al. 2022), during which the UIB community was co-dominated by centric and pennate diatoms, while centric diatoms were associated to development of the bloom in the marginal ice zone and open waters (Lafond et al. 2019; Ribeiro et al. 2022). These varying temporal dynamics within Baffin Bay highlight the differences in bloom phenology linked to the heterogeneity of Arctic outflow shelves (Michel et al. 2015).

We observed an interspecific temporal succession among the planktonic *Thalassiosira* species, *T. nordenskioeldii*, *T. antarctica* and *T. aestivalis/T. pacifica* (Figure 6). *Thalassiosira* species are major contributors to standing stock and carbon export in the Arctic (Arrigo et al. 2012; Krause et al. 2018; Wiedmann et al. 2014). Elemental stoichiometry analysis of *T. nordenskioeldii*, *T. antarctica* and *T. aestivalis* has shown no significant differences in their biovolume, although their carbon quota per biovolume under similar growth conditions varies slightly (Lomas et al. 2019). The impact of the observed temporal succession between these planktonic species on their functional role and carbon export throughout the bloom phases remains therefore to be explored.

Compared to diatoms, pico-sized sympagic and under-ice photosynthetic eukaryotes (*≤* 3 *µ*m) have received less attention (Belevich et al. 2018; Freyria et al. 2021; Lovejoy et al. 2007). Cryptophytes, represented by the genus *Baffinella* and an undescribed Cryptomonadales clade, were among the dominant taxa within the pico-sized sympagic community during the snow-covered stage (Figure 3). Cryptophytes are known for their ability to adapt to a range of light intensities (Greenwold et al. 2019; Heidenreich and Richardson 2020). A time series from the high Arctic Kangerluarsunnguaq fjord (the Danish name for which is Kobbefjord) on the west coast of Greenland, also found cryptophytes and other flagellates as dominant within the sympagic community when light was limited (Mikkelsen et al. 2008). During pre-bloom conditions (stage I), Bolidophyceae (Parmales environmental clade 2) were the dominant picoplanktonic group at all depths (Figure 3). Bolidophyceae have been previously found to be abundant in polar and subarctic regions from DNA sequences and microscope observations (Belevich et al. 2017; Ichinomiya et al. 2016).

The dominance of cryptophytes (and pelagohytes, Figure 3) and bolidophytes within the sympagic and under-ice planktonic communities under low light regimes indicates their important role as primary producers during periods of heavy snow cover (winter and early spring). Early primary producers within the sympagic community like cryptophytes are a rich food source for both ice and early season grazers (Durbin and Casas 2014; Graeve et al. 2001; Kohlbach et al. 2016; Mohan et al. 2016). Although the contribution of Bolidophyceae to the polar food web remains unclear, cells resembling the silicified Bolidophyceae have been reported in fecal pellets of subarctic copepods(Booth et al. 1980; Urban et al. 1993). Additionally, sequences belonging to environmental clade 2 have been recorded from polar-temperate waters (Kuwata et al. 2018), corroborating our biogeography analysis.

During the snow-melt stage, a shift in dominance from bolidophytes to prasinophytes, represented by the genus *Micromonas* (Mamiellophyceae), was observed within the planktonic pico-sized community, except in the surface water layer. The increase in subsurface PAR (Figure 2) combined with a slight increase in water temperatures and in stratification (Oziel et al. 2019), likely favoured the growth of the prasinophyte *Micromonas*. During this period the sea ice started to warm by approximately 1.5 C and the bulk brine volume increased (Oziel et al. 2019). The increase in the relative abundance of cryptophytes, especially *Baffinella*, in the under-ice phytoplankton community in surface water was probably a result of brine flushing of these algae into the water (Figure 3).

Later in the time series, another shift in dominance from cryptophytes to prasinophytes was observed within the sympagic pico-sized community in the 3-10 cm ice layer (Figure 3). A study that reported growth rates of *M. polaris* CCMP2099 and *B. frigidus* CCMP2045 indicated that the prasinophyte has a higher maximum growth rate (0.55 divisions. day*^−^*^1^) than the cryptophyte *Baffinella* (0.40 divisions. day*^−^*^1^) when light was not a limiting factor (Lovejoy et al. 2007). Shorter-term physiological experiments have shown that the optimal growth rates for *Micromonas* were also achieved at a slightly higher temperature range (6*^◦^*C - 8*^◦^*C), than *Baffinella* (4*^◦^*C - 6*^◦^*C) (Daugbjerg et al. 2018; Lovejoy et al. 2007). When light is not limiting, the relatively fast growth of *Micromonas* when compared to *Baffinella* likely confers a strong selective advantage to the prasinophyte, which increases in dominance first within the planktonic community as light becomes more available and later among the sympagic community.

Along with *M. polaris*, *P. pouchetii* also dominated the under-ice bloom pico-size community from the snow-melt stage to the end of the time series (Figure 3). This species is recognized as an important member of the pan-Arctic UIB community during the spring-to-summer transition (Ardyna et al. 2020a; Lasternas and Agustı 2010; Schoemann et al. 2005). It has also been reported in early blooms beneath snow-covered pack ice (Assmy et al. 2017). In our data, a small increase in the abundance of *Phaeocystis* in the nano-size community was observed during the early weeks of the ice-melt stage (Figure 3), which could be due to the formation of cell aggregates (Toullec et al. 2021) or of colonies in the late stages of the spring bloom. Under high-light and low-nutrient conditions, such as those found during the ice-melt stage (Oziel et al. 2019), blooming *Phaeocystis* species tend to form polysaccharide-based mucilaginous colonies that can be millimetres in diameter and serve the functions of energy storage and defence against grazers (Nejstgaard et al. 2007; Schoemann et al. 2005). The *P. pouchetii* ASV obtained during the Green Edge oceanographic campaign was flagged as an indicator of the marginal ice zone and open water sectors within the > 20 *µ*m size fraction, corroborating our results (Ribeiro et al. 2022).

Cell size was another element structuring the sympagic and phytoplanktonic communities (Figure S7). In contrast to the relatively stable environmental conditions of the tropics, seasonality, found in temperate and polar regions, is suggested to prompt regular community reorganization through shifts in species composition, leading to a wider size diversity compared to tropical environments (Acevedo-Trejos et al. 2015).

The considerable amount of novel molecular diversity found in this study emphasizes the need for additional research dedicated to formally describing novel arctic taxa (Figure S6). This is evident even among well-studied groups like diatoms, as illustrated by the fact that most (14) of the taxa observed by SEM lack sequence information and therefore cannot be identified by metabarcoding (Tables S2, S3, Figures S3, S4 and S5). It is noteworthy that among the 1 000 genera catalogued by Fourtanier and Kociolek (1999) and Fourtanier and Kociolek (2003), only 197 have reference 18S rRNA sequence (Guillou et al. 2012). About 50% of the most abundant ASVs in our study have a cultured representative (Figure 5), indicating that culturing methods have been able to capture the diversity among abundant arctic taxa (Balzano et al. 2017; Potvin and Lovejoy 2009; Ribeiro et al. 2020; Supraha et al. 2022). However, this is not the case for the rarer taxa (Figure S6), highlighting the need for renewed culturing efforts and development of isolation methods to fill this gap.

### Biogeography and niche preference

The ongoing poleward flow of warm Atlantic and Pacific waters induces an ecosystem shift in the Arctic Ocean towards a more temperate state, marked by the intrusion of temperate species (Gregory et al. 2019; Oziel et al. 2020; Paulsen et al. 2016). Previous studies have linked taxonomically annotated 18S rRNA sequence “amplicons or barcodes” found in the Arctic with their global occurrence in an effort to detect ongoing shifts within the (phyto)plankton community (Ibarbalz et al. 2023; Supraha et al. 2022). In the present study, the prevailing geographic distributions of most of the ASVs for which it could be allocated (representing 82.5% of the reads in our dataset) were polar (57.5%) and polar-temperate (34%). This contrasts with a previous report by Ibarbalz et al. (2023), which reported, using different approaches (OTU vs. ASV, occurrence vs. indicator species approaches), that although abundant, only 12% of their OTUs were represented by taxa with polar distribution. The high prevalence of polar ASVs in our study may be attributed to two factors. Firstly, Ibarbalz et al. (2023) study included Arctic samples collected in the Atlantic and Pacific inflow shelves, where a high number of non-polar barcodes were detected. In contrast, our sampling site was located on the western coast of Baffin Bay, a region characterized by the outflow of modified Pacific and Arctic waters (Ardyna et al. 2020a). The under-ice water column at the ice camp was dominated by Arctic Water advected southward along Baffin Island (Oziel et al. 2019). Secondly, in contrast to the study of Ibarbalz et al. (2023), the presence of sympagic communities and taxa present at the transition between the late winter and early spring in our dataset might have contributed to an increased number of polar barcodes.

Most sympagic ASVs were classified as polar. Some exceptions (39 of 306) corresponded to ASVs found at latitudes below the polar circle from the Baltic Sea and Hudson Bay, which are seasonally ice-covered (European Union-Copernicus Marine Service 2021). The sympagic community also exhibited more ASVs significantly associated with ice as a substrate than water (Figure 5). These ASVs may represent sea ice specialists which can cope with the strong gradients of salinity, temperature and light found in the ice (Arrigo 2014). For example, some cryptophyte species like *B. frigidus* have been characterized as euryhaline, growing at salinities ranging from 5 to 35 (Daugbjerg et al. 2018). The *B. frigidus* ASV_c699cb1809 and ASV_f8124a0c4a were 100% similar to the sequences obtained from the strains CCMP2045 (GQ375264) and RCC5289 (OR736128), isolated from Baffin Bay waters (Daugbjerg et al. 2018) and sea ice, respectively (Ribeiro et al. 2020). The latter ASV (ASV_f8124a0c4a) was flagged as an indicator of ice (Figure 5A) and showed a restricted polar distribution (Figure 6) which suggests that it probably represents an ice ecotype. Additional studies addressing the intraspecific variability in growth optima between ice and water isolates are required for an in-depth characterization of the niche preferences and fitness of the cryptophyte genus *Baffinella*.

Another example of potential ice specialist taxa is represented by the polar ASV_58a35e6970 which was classified as undescribed Pelagomonadales clade A1, also flagged as an indicator of the ice community (Figure 5A) and 100% similar to the sequence obtained from the Arctic pelagophyte CCMP2097 (EU247837), a strain isolated from sea ice. Metatranscriptomic analysis has revealed that CCMP2097 possesses specific adaptations to cold saline conditions, such as those found in sea ice micro-environments. This includes differential expression of several antifreeze proteins, an ice-binding protein, and an acyl-esterase involved in cold adaptation (Freyria et al. 2022).

As the bloom developed, we observed a transition from polar to polar-temperate ASVs in the smallest size fraction, while polar ASVs represented by undescribed groups of pennate (Bacillariophyceae) and centric (Mediophyceae) diatoms were major contributors to the micro-sized planktonic community across all bloom stages and nearly all depths (Figure 3). The predominance of polar ASVs within the micro-sized planktonic communities might be explained by their lower dispersal rate compared to the pico- and nano-sized taxa. Body size plays an important role in determining spatial patterns for planktonic organisms, where limitation of dispersal is expected to increase with body size (Villarino et al. 2018). Although diatoms associated with sea ice in the Arctic have been reported as endemic taxa (Poulin et al. 2011; von Quillfeldt 2000), our results corroborate the idea that Arctic endemism is also relatively common among planktonic diatoms, especially within the micro-plankton where a smaller connection between distant diatoms communities is expected (Supraha et al. 2022).

A few ASVs, in low abundance, were allocated as temperate (6) and cosmopolitan (11), some representing taxa with broad biogeographical signatures (Figure 4). The cosmopolitan *Phaeocystis* ASV_8050f737e4 was 100% similar to sequences obtained from the species *P. jahnii* which was initially described from the Mediterranean Sea (Zingone et al. 1999) and reported in the warmer waters of the southeastern East China Sea (Song et al. 2023). The ecological versatility (i.e., broad geographic distribution with species found across several gradients of temperatures and nutrient conditions) of this genus stemming from their ability to grow mixotrophically (Koppelle et al. 2022; Mars Brisbin et al. 2022) aligns well with suggestions of a shift towards *Phaeocystis*-dominated blooms in future Arctic scenarios, which will have implications for phytoplankton community structure and trophic energy transfer (Lasternas and Agustı 2010; Rokitta et al. 2023). The detection of the warm water species *Phaeocystis jahnii* ASV_8050f737e4 in our dataset might be an indicator of the ongoing turnover of the phytoplankton community in the Arctic outflow shelves.

While barcodes found in the Arctic with non-polar occurrence in global datasets may represent indicators of ongoing species displacement within the Arctic planktonic community, they also stress the limitation of using barcode sequences, such as the variable regions of 18S rRNA, when describing biogeographic patterns of plankton species. The *B. prasinos* ASV_4580ad6202 was allocated as cosmopolitan and although the genus *Bathycoccus* is considered cosmopolitan, the analysis of metagenomes (Vaulot et al. 2012), of nuclear genomes (Dennu et al. 2023; Vannier et al. 2016) and of the internal transcribed spacer 2 (Bachy et al. 2021) have suggested the existence of distinct *Bathycoccus* ecotypes and species, including a polar genotype (Dennu et al. 2023), all of which have identical 18S rRNA sequences and therefore cannot be discriminated by metabarcodes from the V4 or V9 regions of the 18S rRNA gene. Among diatoms, the polar-temperate ASV_2976201d4c represents at least two distinct *Thalassiosira* species, *T. pacifica* and *T. aestivalis*, which share identical 18S rRNA V4 regions and can be only distinguished by scanning electron microscopy (i.e, *T. pacifica* was detected by SEM in our samples while *T. aestivalis* was absent). In the future, the combined use of more resolutive taxonomic markers, such as ITS or long-read amplicons capable of distinguishing cryptic genotypes will likely unveil additional biogeographic patterns.

Finally, a large proportion of ASVs (53%), especially originating from the sympagic community, remained unallocated, probably due to the lack of sufficient metabarcoding datasets from ice samples (Figure S8) in the metaPR2 database (Vaulot et al. 2022). Furthermore, due to the limited number of datasets available from the Antarctic, we have opted for a more conservative approach by not separating the polar biogeographical categories into Arctic and Antarctic categories. Some of our barcodes may indeed represent bipolar taxa (Segawa et al. 2018; Wolf et al. 2015), hence we avoided unsubstantiated claims of “Arctic-exclusive” barcodes or taxa despite their strong occurence in the Arctic (Figure 4).

## Conclusion

The current study offers evidence that the classes Pelagophyceae, Bolidophyceae and Cryptophyceae may play a more important role within the pico-sized communities in and under the ice pack during pre-bloom conditions than previously thought. The significant prevalence of undescribed biodiversity (both undescribed and uncultivated taxa) in this study, even among morphologically well-described groups such as diatoms, underscores the urgent need for more research focusing on formally describing new taxa. Additionally, novel isolation methods and efforts are necessary to further elucidate the taxonomic and functional biodiversity of Arctic microbial eukaryotes. The report that *B. frigidus* RCC5289, a strain isolated during Green Edge from ice samples, likely represents an ice ecotype is an illustration of the importance of microdiversity analyses. The use of metabarcoding data from 2,874 worldwide marine samples offers a valuable approach for developing monitoring strategies of microbial eukaryotes in a changing Arctic Ocean. As the number of metabarcoding studies increases exponentially (Lopes dos Santos et al. 2022), such analyses will become more robust and statistically significant.

## Acknowledgements

We are grateful to Michel Poulin (Canadian Museum of Nature) and Józef Wiktor (Institute of Oceanology of the Polish Academy of Sciences) for their assistance in the identification of diatoms in the SEM samples. This project would not have been possible without the support of the hamlet of Qikiqtarjuaq and the members of the community, as well as the Inuksuit School and its Principal, Jacqueline Arsenault. The project was conducted under the scientific coordination of the Canada Excellence Research Chair in Remote Sensing of Canada’s new Arctic frontier and the CNRS and Université Laval Takuvik Joint International laboratory (UMI3376). The success of the field campaign is attributed to the contribution of Andrew Wells, Maxime Benoît-Gagné and Emmanuel Devred from the Takuvik laboratory, as well as Robert Hodgson from the University of Manitoba. We also thank Québec-Océan and the Polar Continental Shelf Program for their in-kind contribution in terms of polar logistics and scientific equipment. We are grateful to the ABIMS platform of the FR2424 (CNRS, Sorbonne Université) for providing excellent computer resources.

## Author contributions statement

ALS and DV have contributed to study conception and design. ALS, PG, IP and DV have acquired the samples. ALS, CGR, FL, PG, and IP have processed the samples and produced data. CS, ALS, DV, CL, and CGR have analysed and interpreted the data. CS and ALS have drafted the manuscript. All authors have revised and approved the final manuscript.

## Data availability

Raw Illumina sequences were deposited to GenBank under project PRJNA810431. All codes and data used in this study can be found in https://github.com/clarencesimple/SIM_GreenEdge_IceCamp/tree/main

## Supplementary material

**Table S1:** Number of samples obtained from metaPR^2^ (Dataset version 2.0) used for biogeographical distribution analysis

| Region | No. of samples |
| --- | --- |
| Arctic | 486 |
| North temperate | 1315 |
| Tropical | 196 |
| South temperate | 743 |
| Antarctic | 134 |

**Table S2:**
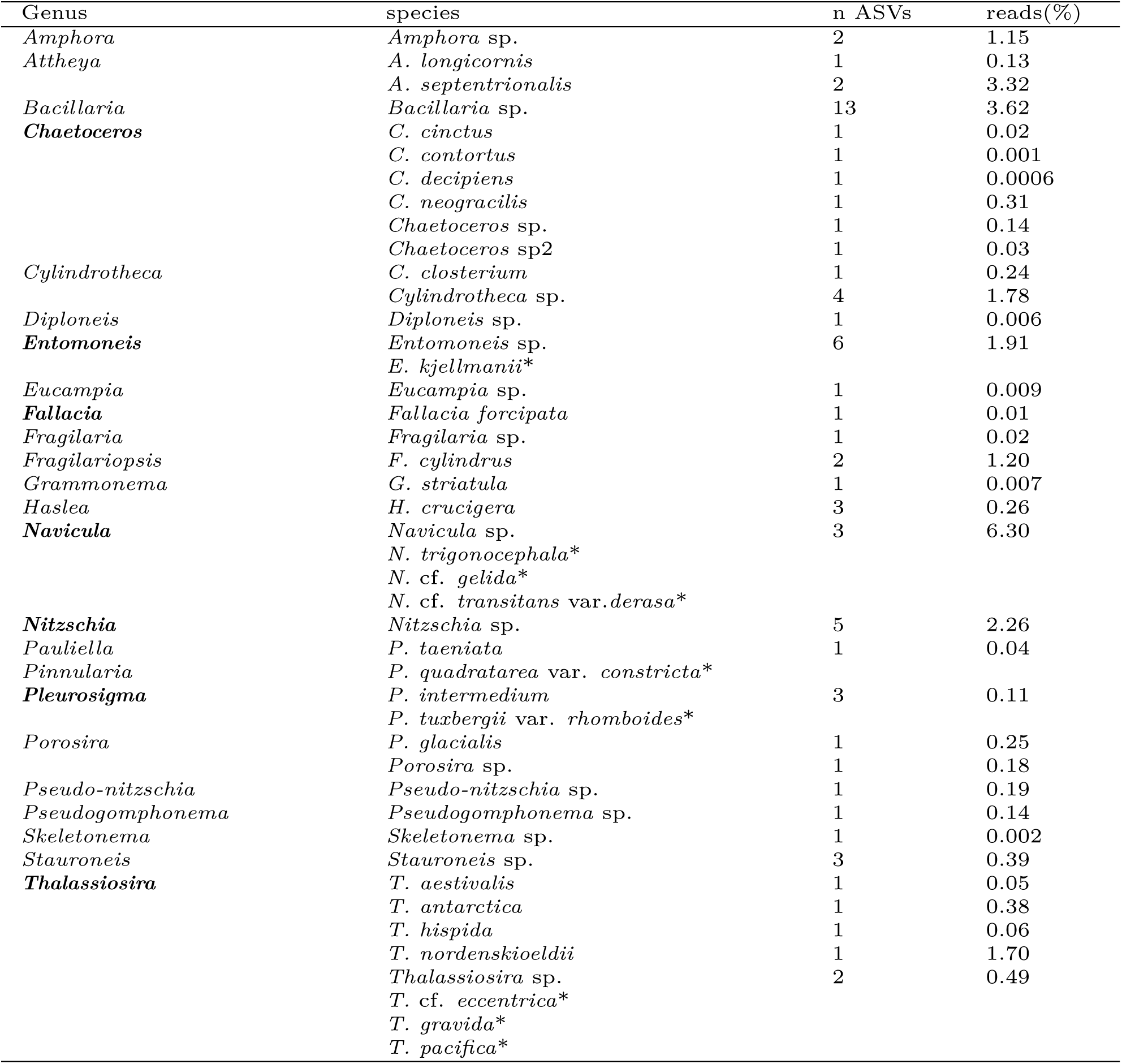
Sympagic diatom taxa identified by metabarcoding and SEM (Figure S3). Genera and species with *≥* 80 bootstrap sequence support are listed. Percentage reads corresponds to the sum of DNA reads from taxon in ice over total photosynthetic DNA reads in ice. Genus names in bold have been identified by both methods. * Species only identified by SEM

**Table S3:** Phytoplanktonic diatom taxa identified by metabarcoding and SEM (Figures S4 and S5). Genera and species with *≥* 80 bootstrap sequence support are listed. Percentage reads corresponds to the sum of DNA reads from taxa in water over total photosynthetic DNA reads in water. Genus and species names in bold have been identified by both methods. * Species only identified by SEM

| Genus | species | n ASVs | reads(%) |
| --- | --- | --- | --- |
| *Achnanthes* * | *Achnanthes* sp.* |  |  |
| *Amphora* | *Amphora* sp. | 2 | 0.30 |
| ***Attheya*** | *A. longicornis* | 1 | 0.005 |
|  | ***A. septentrionalis*** | 1 | 0.84 |
| *Bacillaria* | *Bacillaria* sp. | 10 | 0.73 |
| *Chaetoceros* | *C. cinctus* | 1 | 0.05 |
|  | *C. contortus* | 1 | 0.05 |
|  | *C. debilis* 1 | 1 | 0.01 |
|  | *C. decipiens* | 1 | 0.001 |
|  | *C. neogracilis* | 1 | 1.12 |
|  | *Chaetoceros* sp. | 7 | 2.55 |
|  | *Chaetoceros* sp2 | 1 | 0.06 |
| *Cocconeis* * | *Cocconeis* sp.* |  |  |
| *Cylindrotheca* | *C. closterium* | 1 | 0.03 |
|  | *Cylindrotheca* sp. | 3 | 0.62 |
| *Detonula* | *D. confervacea* | 1 | 0.01 |
| ***Entomoneis*** | *Entomoneis* sp. | 5 | 0.43 |
|  | *E. kjellmanii* var. *kariana* * |  |  |
| *Eucampia* | *Eucampia* sp. | 1 | 0.005 |
| *Fallacia* | *F. forcipata* | 1 | 0.005 |
| *Fragilaria* | *Fragilaria* sp. | 4 | 3.84 |
| *Fragilariopsis* | *F. cylindrus* | 2 | 6.03 |
| *Grammonema* | *G. striatula* | 1 | 0.003 |
| *Haslea* | *H. crucigera* | 2 | 0.11 |
| *Licmophora* | *L. gracilis* | 1 | 0.001 |
| *Mediolabrus* * | *Mediolabrus comicus* * |  |  |
| ***Navicula*** | *Navicula* sp. | 3 | 1.65 |
|  | *N. superba* * |  |  |
|  | *N. cf. pagophila* var. *manitounukensis* * |  |  |
|  | *N. trigonocephala* var. *depressa* * |  |  |
| *Nitzschia* | *Nitzschia* sp. | 4 | 1.33 |
| *Pauliella* | *P. taeniata* | 1 | 0.66 |
| *Pinnularia* * | *Pinnularia quadratarea* var. *constricta* * |  |  |
| ***Pleurosigma*** | *P. intermedium* | 3 | 0.15 |
|  | *P. stuxbergii* var. *rhomboides* * |  |  |
| ***Porosira*** | *P. glacialis* | 1 | 1.53 |
|  | *Porosira* sp. | 1 | 1.94 |
| ***Pseudo-nitzschia*** | *Pseudo-nitzschia* sp. | 3 | 8.17 |
|  | cf. *Pseudo-nitzschia obtusa* * |  |  |
| ***Pseudogomphonema*** | *Pseudogomphonema* sp. | 1 | 0.11 |
|  | *P. arcticum* * |  |  |
|  | *P. septentrionale* var. *angustatum* * |  |  |
| *Skeletonema* | *S. marinoi* | 1 | 0.004 |
|  | *Skeletonema* sp. | 1 | 0.13 |
| *Stauroneis* | *Stauroneis* sp. | 2 | 0.17 |
| ***Thalassiosira*** | *T. aestivalis* | 1 | 0.93 |
|  | *T. anguste-lineata* | 1 | 0.02 |
|  | *T. antarctica* | 1 | 2.93 |
|  | ***T. gravida*** | 1 | 0.43 |
|  | *T. hispida* | 1 | 0.12 |
|  | ***T. nordenskiöldii*** | 1 | 1.93 |
|  | *Thalassiosira* sp. | 3 | 2.92 |
|  | *T. cf. eccentrica* * |  |  |
|  | *T. hyalina* * |  |  |

**Table S4:** ANOSIM R and p values obtained by comparing samples clustered based on size fractions, substrate, and bloom stages

|  | **R-value** | **p-value** |
| --- | --- | --- |
| **All samples** |  |  |
| Size fraction (pico, nano, micro) | 0.558 | 0.001 |
| Substrate (ice:water) | 0.460 | 0.001 |
| **Ice samples** |  |  |
| Size fraction (pico, nano, micro) | 0.671 | 0.001 |
| Bloom stages (I:II:III) | 0.173 | 0.004 |
| **Water Samples** |  |  |
| Size fraction (pico, nano, micro) | 0.726 | 0.001 |
| Bloom stages (I:II:III) | 0.273 | 0.001 |

**Figure S1.**
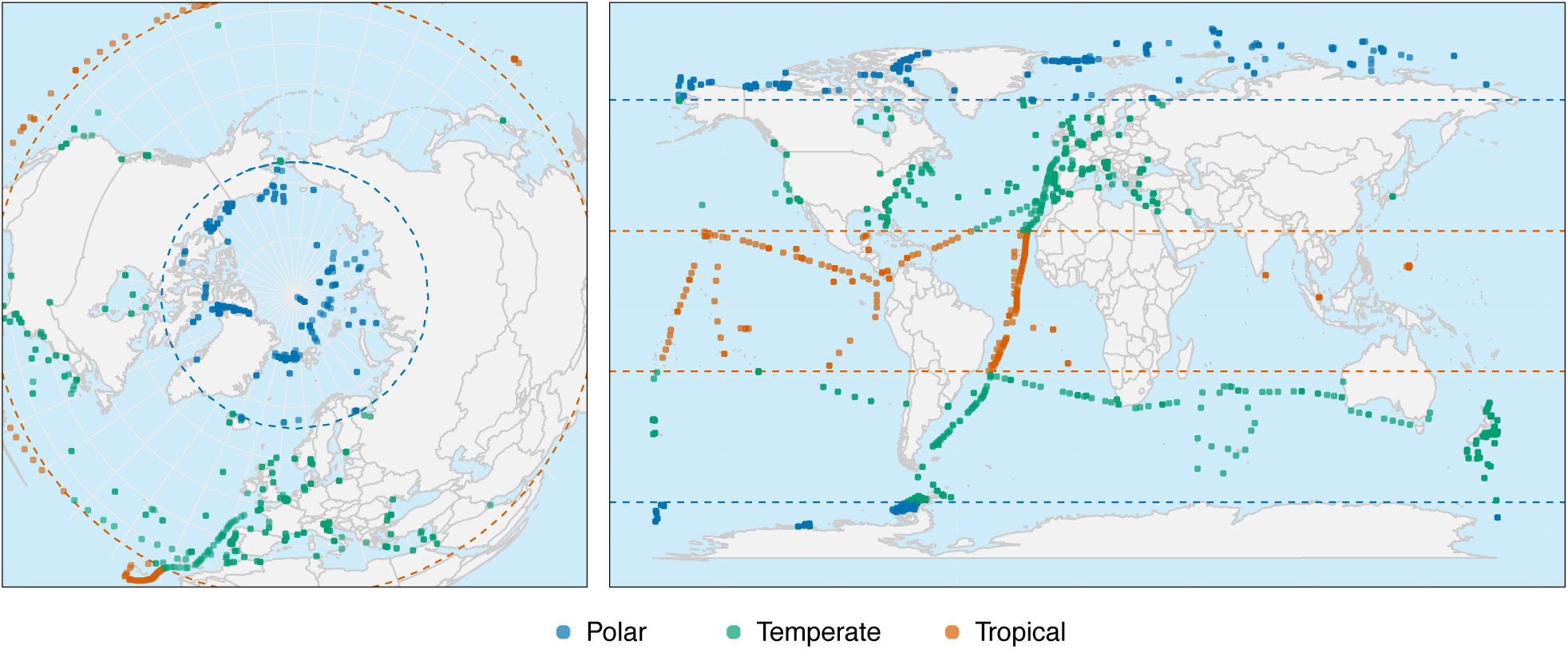
Pan-Arctic (left) and global (right) distribution of coastal and oceanic samples from metaPR^2 d^atabase (Datasets version: 2.0) used for biogeographical analyse^s^in this study.

**Figure S2.**
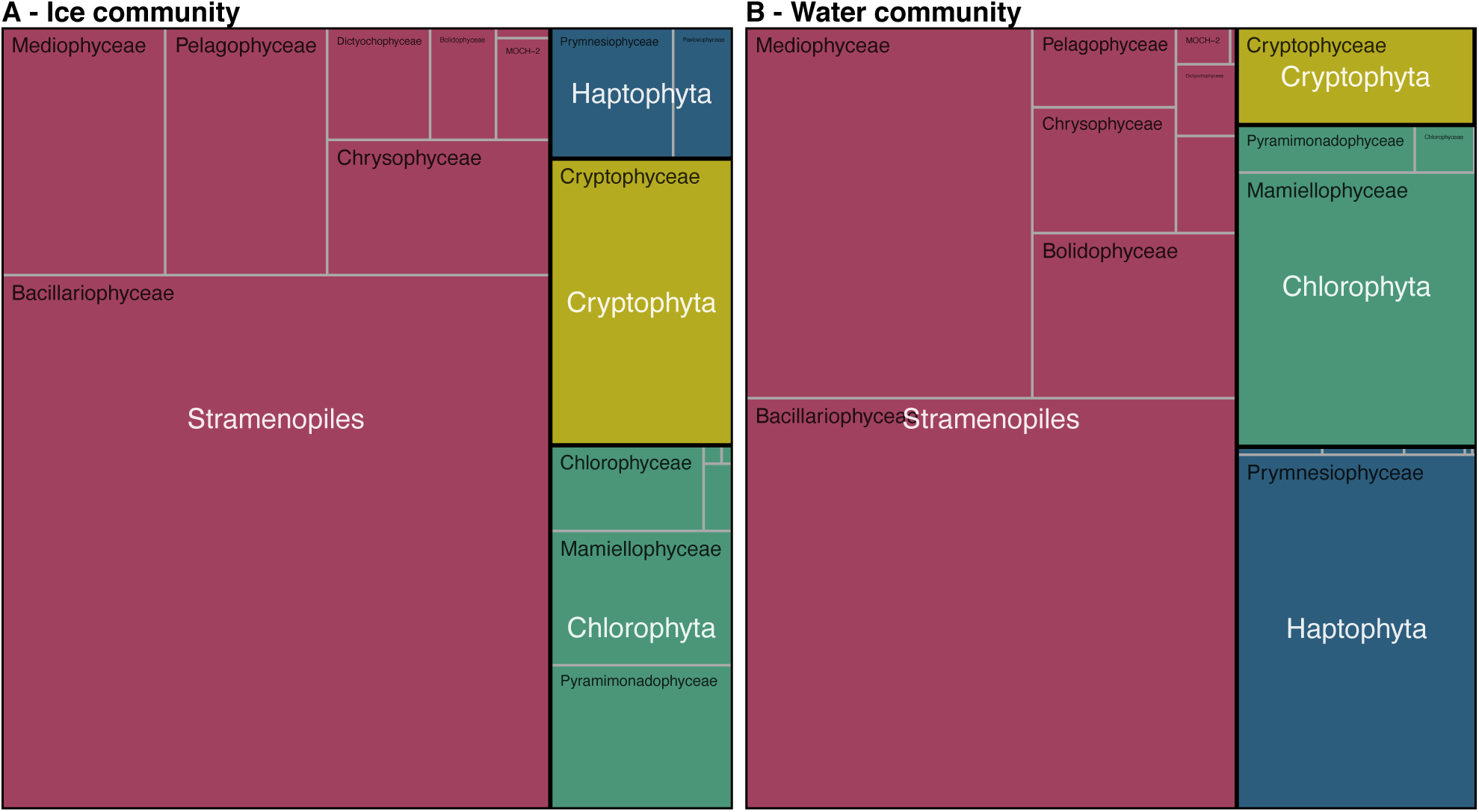
Community composition from most abundant photosynthetic taxa at class level from (A) ice and (B) water samples. Proportional area charts of normalised abundance of 18S V4 reads based on division (white labels) and class (black labels).

**Figure S3.**
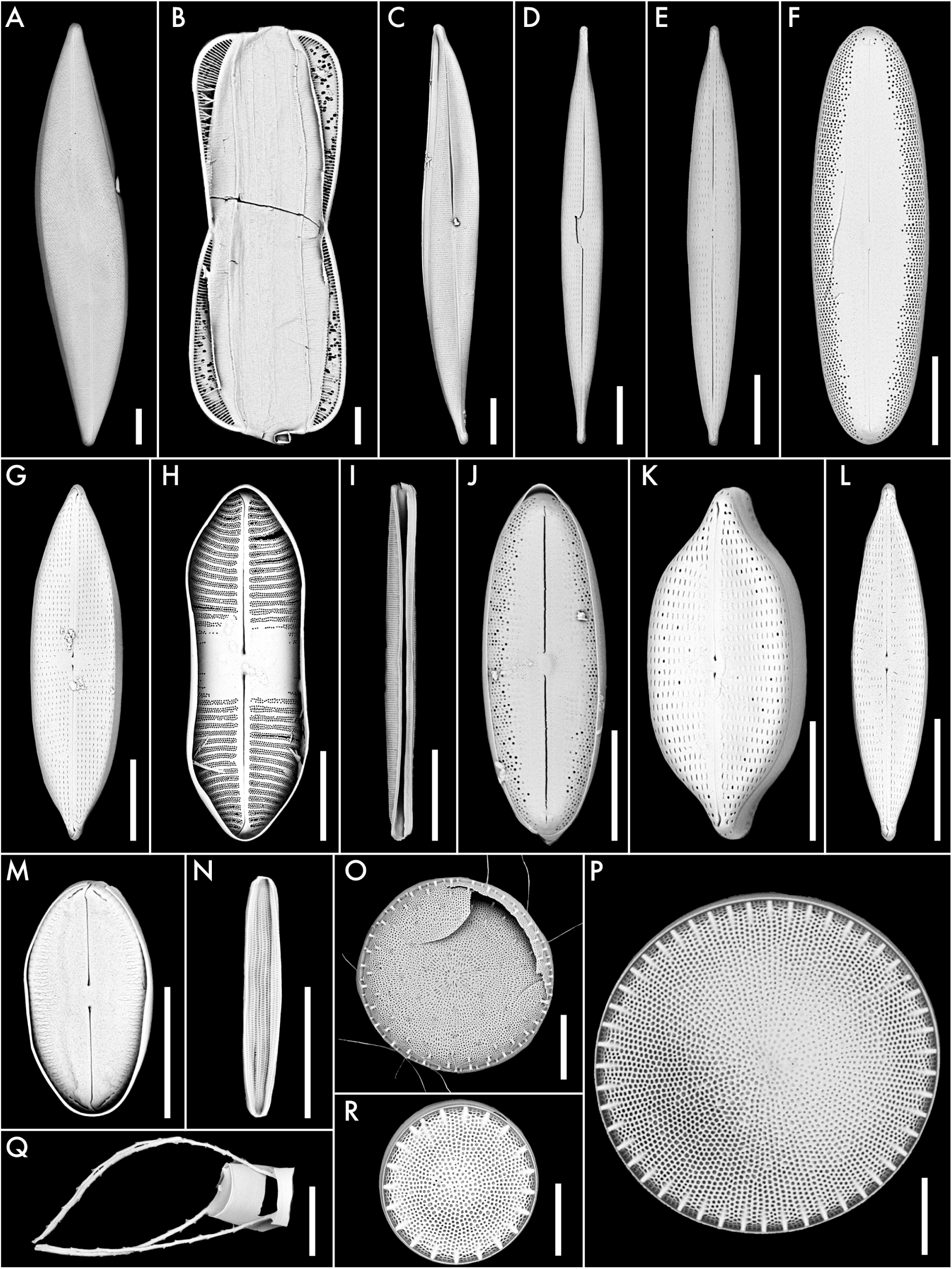
Diversity of diatoms in ice samples from scanning electron microscopy. Size bars correspond to 10 *µ*m. A) *Pleurosigma stuxbergii* var. *rhomboides*; B) *Entomoneis kjellmanii*; C) *Gyrosigma concilians*; D) *Navicula* sp.; E) *Navicula* cf. *directa*; F) Raphid pennate G) *Navicula* cf. *gelida*; H) *Pinnularia quadratarea* var. *constricta*; I) *Nitzschia* sp.; J) cf. *Fallacia* sp.; K) *Navicula trigonocephala*; L) *Navicula transitans* var. *derasa*; M) *Fallacia* sp.; N) *Amphora* sp.; O) *Thalassiosira gravida*; P) *Thalassiosira pacifica*; Q) *Chaetoceros* sp.; R) *Thalassiosira* cf. *eccentrica*.

**Figure S4.**
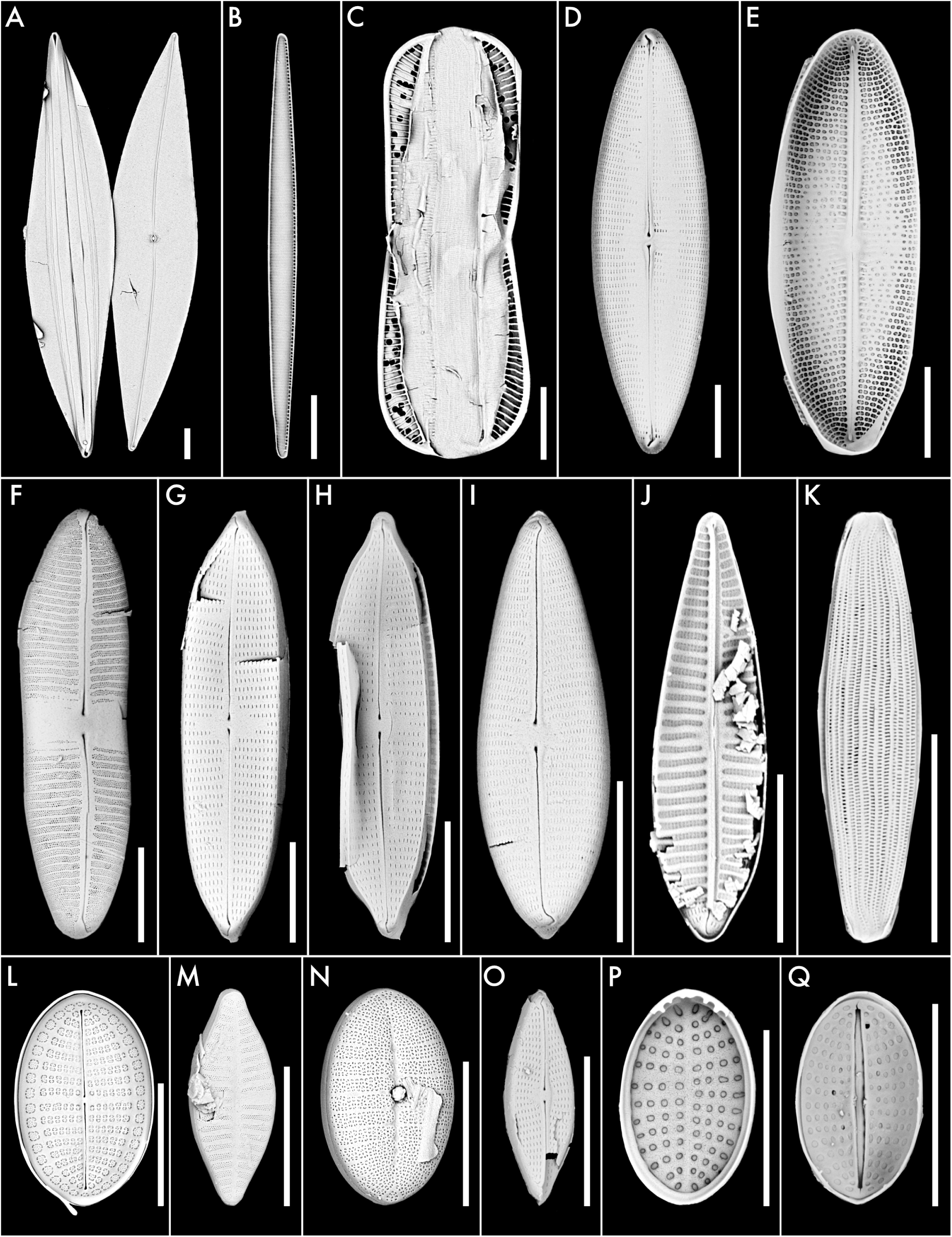
Diversity of pennate diatoms in water samples from scanning electron microscopy. Size bars correspond to 10 *µ*m. A) *Pleurosigma stuxbergii* var. *rhomboides*; B) *Pseudo-nitzschia* sp.; C) *Entomoneis kjellmanii* var. *kariana*; D) *Navicula* sp.; E) *Navicula* cf. *pagophila* var. *manitounukensis*; F) *Pinnularia quadratarea* var. *constricta*; G) *Navicula trigonocephala* var. *depressa*; H) *Navicula trigonocephala* var. *depressa*; I) *Pseudogomphonemma arcticum*; J) *Pseudogomphonema septentrionale* var. *angustatum*; K) *Amphora* sp.; L) *Cocconeis* sp.; M) *Achnanthes* sp.; N) *Cocconeis* sp.; O) *Navicula* sp.; P) *Cocconeis* sp.; Q) *Cocconeis* sp.

**Figure S5.**
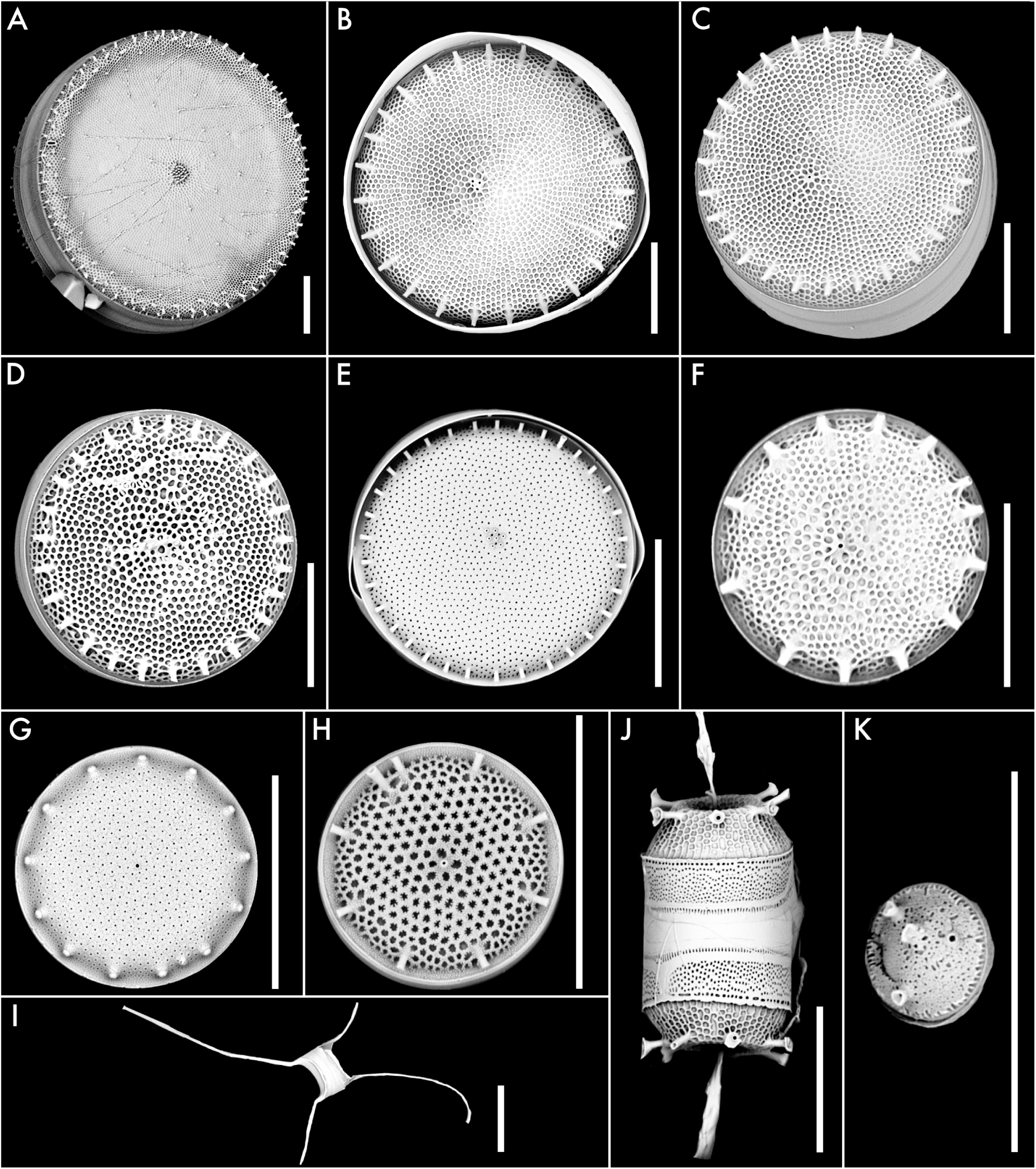
Diversity of centric diatoms in water samples from scanning electron microscopy. Size bars correspond to 10 *µ*m. A) *Thalassiosira gravida*; B) *Thalassiosira* sp.; C) *Thalassiosira* cf. *eccentrica*; D) *Thalassiosira* sp.; E) *Thalassiosira hyalina*; F) *Thalsssiosira* sp.; G) *Thalassiosira* sp.; H) *Thalassiosira* sp.; I) *Attheya septentrionalis*; J) *Thalassiosira nordenskioeldii*; K) *Mediolabrus comicus*.

**Figure S6.**
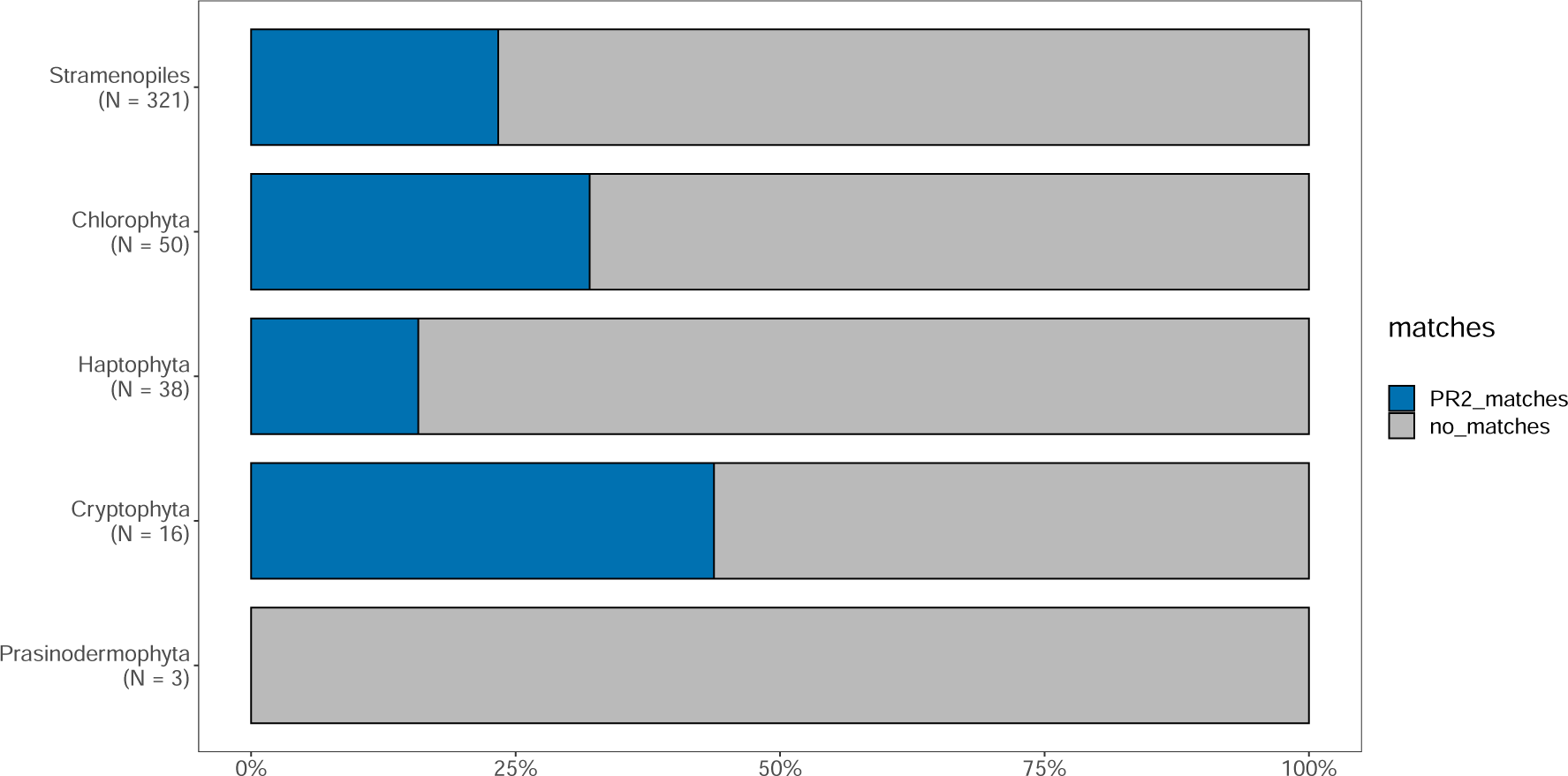
Proportion of ASVs within each division with 100% match to PR^2 s^equences fro^m^ cultures

**Figure S7.**
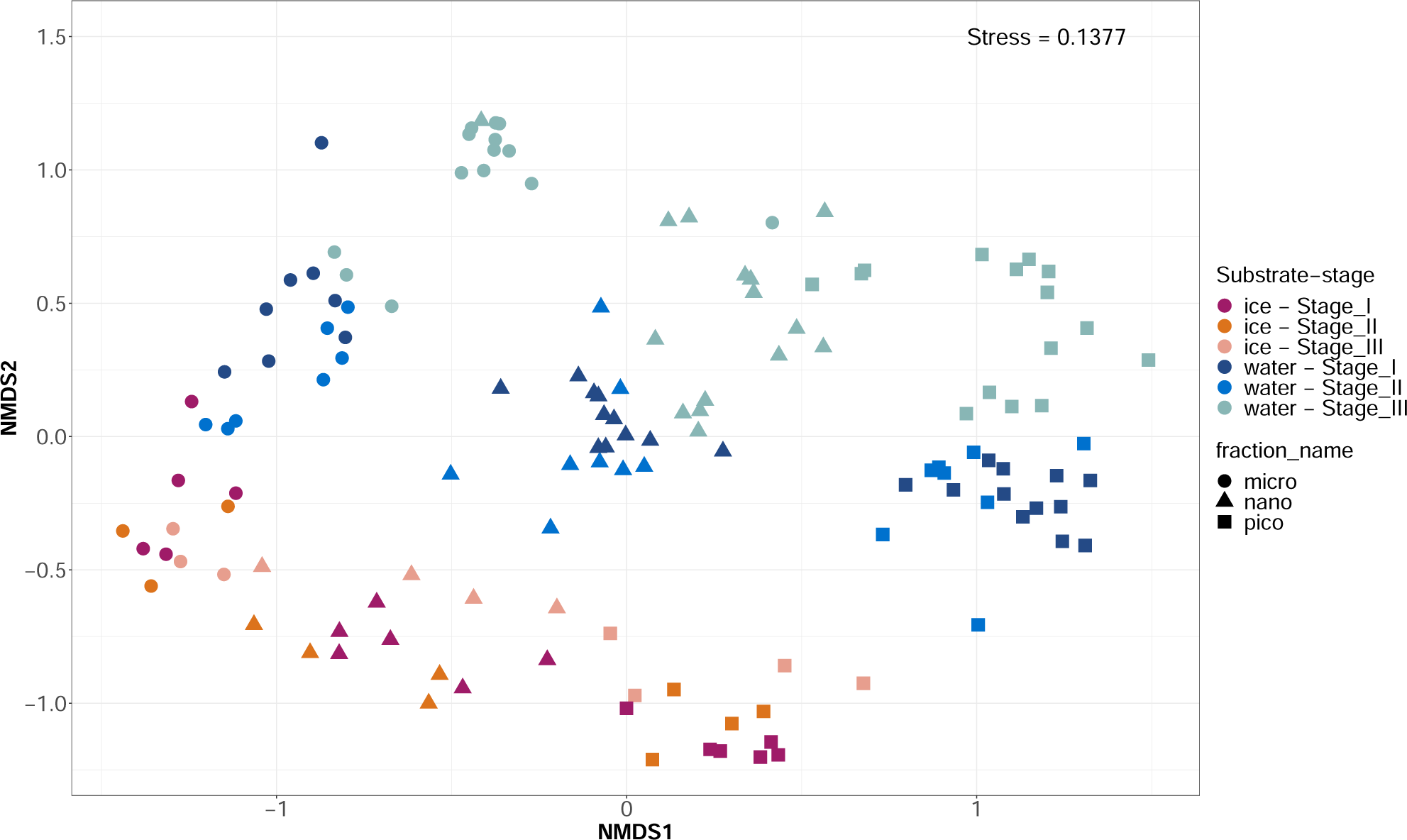
Non-metric multidimensional scaling (NMDS) analysis based on Bray-Curtis dissimilarities of the photosynthetic community composition at ASV level. Labels according to substrate-stage and size fraction.

**Figure S8.**
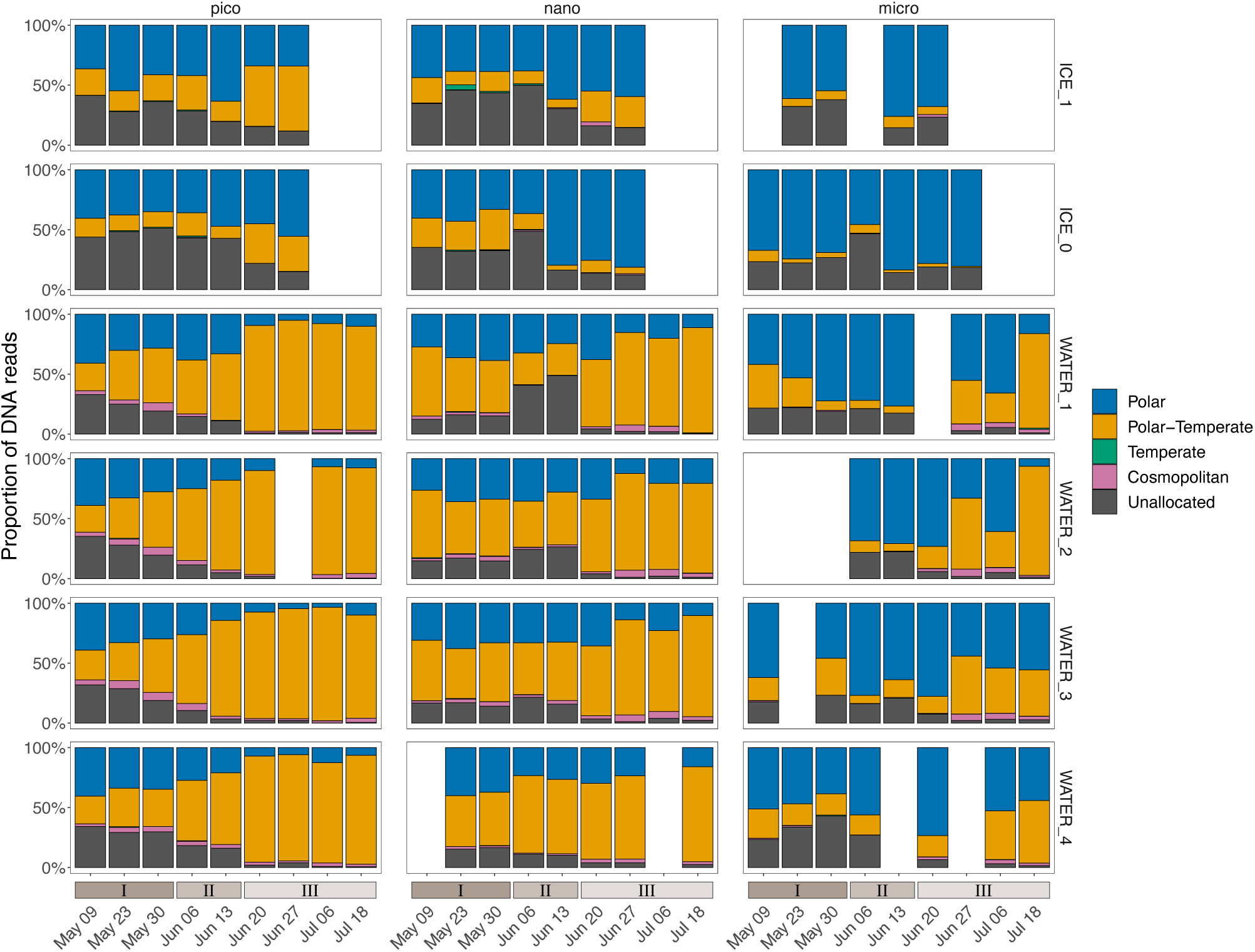
Temporal variation of ASVs according to their biogeographical distribution for each size fraction and each sampled layer.

**Figure S9.**
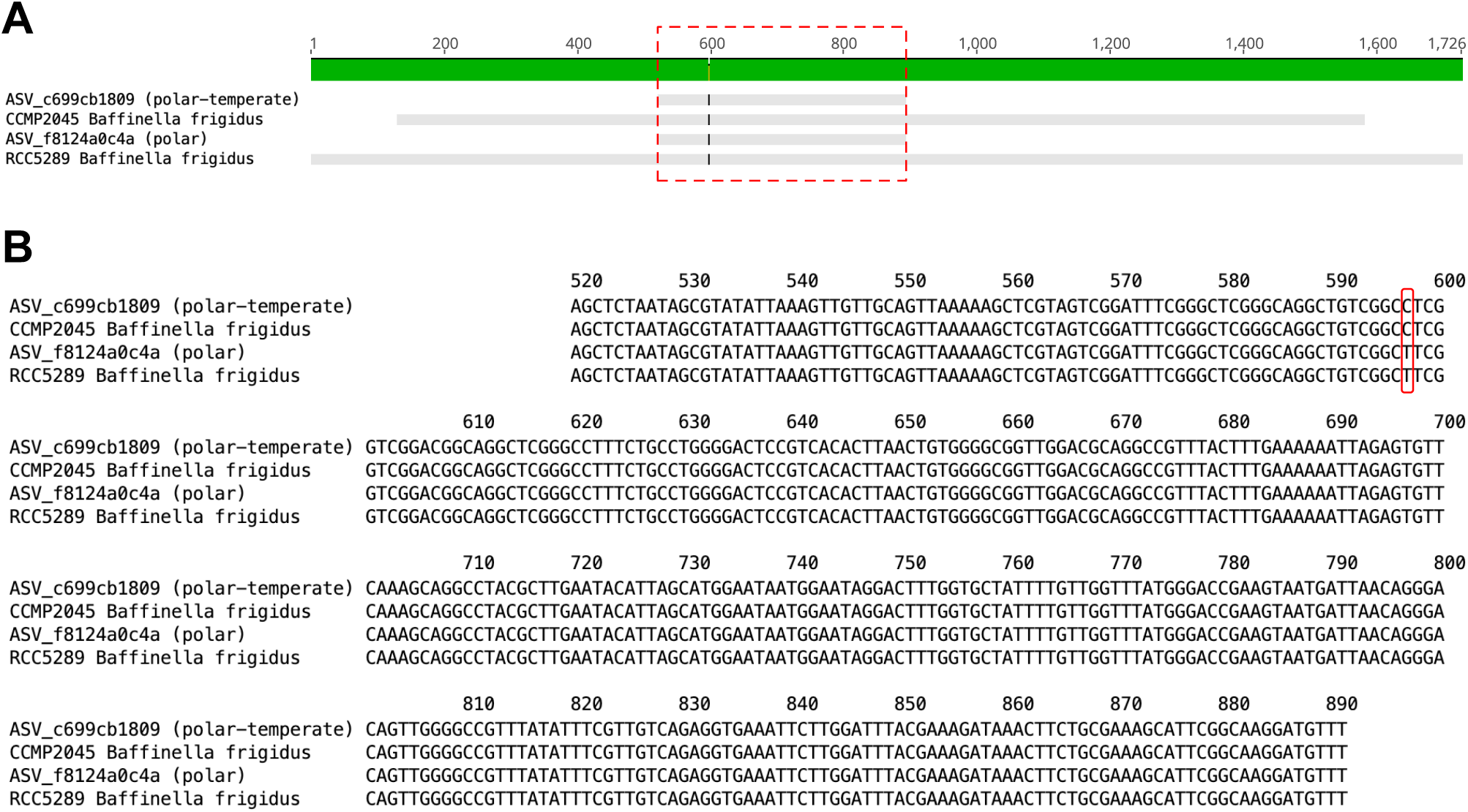
(A) DNA sequence alignment of two 18S V4 metabarcoding ASVs assigned to Baffinella frigidus, and the two partial 18S sequences from cultures. (B) Sequences from the red box showing single nucleotide differences at position 597. ASV_c699cb1809 has 100% similarity with CCMP2045 while ASV_f8124a0c4a has 100% similarity with RCC5289.

